# Multiscale model of the physiological control of myocardial perfusion to delineate putative metabolic feedback mechanisms

**DOI:** 10.1101/2021.08.04.455088

**Authors:** Hamidreza Gharahi, C. Alberto Figueroa, Johnathan D. Tune, Daniel A. Beard

**Affiliations:** Section of Vascular Surgery, Department of Surgery, University of Michigan, Ann Arbor, MI; Department of Molecular and Integrative Physiology, University of Michigan, Ann Arbor, MI; Department of Biomedical Engineering, University of Michigan, Ann Arbor, MI; Department of Physiology and Anatomy, University of North Texas Health Sciences Center, Fort Worth, TX

## Abstract

Coronary blood flow is tightly regulated to ensure that myocardial oxygen delivery meets local metabolic demand via the concurrent action of myogenic, neural, and metabolic mechanisms. While several competing hypotheses exist, the specific nature of the local metabolic mechanism(s) remains poorly defined. To gain insights into the viability of putative metabolic feedback mechanisms and into the coordinated action of parallel regulatory mechanisms, we applied a multi-scale modeling framework to analyze experimental data on coronary pressure, flow, and myocardial oxygen delivery in the porcine heart in vivo. The modeling framework integrates a previously established lumped-parameter model of myocardial perfusion used to account for transmural hemodynamic variations and a simple vessel mechanics model used to simulate the vascular tone in each of three myocardial layers. Vascular tone in the resistance vessel mechanics model is governed by input stimuli from the myogenic, metabolic, and autonomic control mechanisms. Seven competing formulations of the metabolic feedback mechanism are implemented in the modeling framework, and associated model simulations are compared to experimental data on coronary pressures and flows under a range of experimental conditions designed to interrogate the governing control mechanisms. Analysis identifies a maximally likely metabolic mechanism among the seven tested models, in which production of a metabolic signaling factor is proportional to MVO_2_ and delivery proportional to flow. Finally, the identified model is validated based on comparisons of simulations to data on the myocardial perfusion response to conscious exercise that were not used for model identification.

## Introduction

The left ventricular myocardium extracts ~70-80% of the oxygen delivered from the arterial blood under normal resting conditions. A consequence of this high degree of myocardial oxygen extraction is that increases in myocardial oxygen demand (such as during exercise) require proportionate increases in perfusion to the myocardium to maintain adequate tissue oxygenation. This tight coupling of myocardial oxygen delivery with metabolism is observed to occur not only during physiologic increases in myocardial oxygen consumption (MVO_2_), but also following perturbations to coronary perfusion pressure (CPP), which are crucial, especially in response to proximal stenotic lesions of epicardial coronary arteries.

Our understanding of the mechanisms underlying the balance between coronary blood flow and MVO_2_ remains limited. Putatively, coronary flow regulation is mediated via the concurrent action of three major control mechanisms: (1.) a myogenic mechanism where resistance arterioles constrict in response to increases in vascular wall tension; (2.) an autonomic mechanism were stimulation of β-receptors on arterial smooth muscle leads to vasodilation; and (3.) a metabolic mechanism where an increase in MVO_2_ promotes local vasodilation. These mechanisms work in parallel and converge on common end-effector pathways of coronary microvascular resistance; i.e. vascular smooth muscle tone. Moreover, the actions of these control mechanisms are influenced by structural and contractile properties of the myocardium and vary transmurally across layers of the myocardium. Precisely how these purported mechanisms work together dynamically and regionally, in the beating heart where cardiac contraction causes a constricting force on the vessels in the walls of the heart, impeding flow, particularly in the subendocardium, during systole merits further investigation. Furthermore, while several competing hypotheses exist, the specific nature of local metabolic mechanism(s) remains poorly defined.

The paradigm of local metabolic control of coronary blood flow centers around the hypothesis that vasoactive metabolites are produced in proportion to the prevailing level of myocardial oxygenation. This hypothesis is attractive in that it provides a mechanism directly linking increases in metabolism and reductions in tissue oxygen tension (typically index by coronary venous PO_2_) with commensurate changes in coronary blood flow [1]. However, this general framework offers little insight into if or how specific metabolites (e.g., adenosine and NO) or pathways (e.g., end-effector K+ channels) contribute to myocardial oxygen supply/demand balance. In addition, prior observations that coronary venous PO_2_ does not directly correlate with changes coronary blood flow during increases in MVO_2_ [2, 3] or perturbations in CPP [4] fail to support the metabolic hypothesis as proposed. Thus both the general framework of as well as the specific molecular signals involved in the local metabolic control of coronary blood flow continues to linger as one of, if not the most, highly contested mysteries of the coronary circulation to this day.

The goal of this study is to address these knowledge deficits through model-based analysis of experimental data on in vivo coronary flow regulation in response to changes in perfusion pressure, MVO_2_, and the oxygen carrying capacity of the blood. Specifically, a multi-scale model of coronary flow regulation was constructed to account for putative myogenic, neural, and metabolic mechanisms as well as transmural dynamics of blood circulation and ventricular-vascular interactions. Model simulations were compared to data on coronary flow and zero-flow coronary pressure recordings obtained distal to epicardial occlusions *in vivo* in pigs [4] and subendocardial-to-subepicardial flow ratio. Coronary flow and zero-flow pressure transients were obtained over a range of initial perfusion pressures and under control conditions as well as during hemodilution and hemodilution with dobutamine infusion.

The model used to analyze these data is built based on the lumped three-layer model of myocardial perfusion [5] and the myogenic autoregulatory vessel model of Carlson and Secomb [6], with additional components representing β-mediated vasodilation and a metabolic feedback mechanism. Several competing representations of the metabolic mechanism were implemented and tested against the data. Using maximal-likelihood structural and parametric model identification to rule out and refine hypotheses, we identify a novel formulation of the metabolic mechanism in which the metabolic signal is represented as proportional to metabolic rate and flow. This mechanism can be interrupted as representing an *ephemeral metabolic signal* for which production is proportional to MVO_2_ and delivery proportional to flow. Potential candidates for such a signal include short-lived reactive species such as H_2_O_2_ [7]. The identified model is further validated based on comparisons of simulations to data on the myocardial perfusion response to conscious exercise that were not used for model identification.

## Methods

### Experimental Data for Model Identification and Validation

Experimental data for model identification were obtained from prior studies on coronary blood flow autoregulation, including measurements of distal pressure transients in the left-anterior descending (LAD) arterial tree following proximal occlusion of the main LAD trunk in anesthetized pigs. The experimental details are described in Kiel et al. [4]. In brief, for a given experimental condition, the LAD was initially perfused at a constant baseline pressure by a servo-controlled pump. After a steady baseline flow at a given perfusion pressure was attained, the LAD trunk was occluded and the resulting decay in arterial pressure at a distal epicardial location was measured. The aortic pressure was continuously measured throughout the experiment via a femoral artery catheter.

Example pressure transients obtained under control conditions (without dobutamine infusion or hemodilution) are illustrated in Figure 1A. In these data sets each of the six sub-panels in panel A corresponds to a different average initial perfusion pressure, ranging from 40 to 140 mmHg. The perfusion pressure oscillated around the set CPP level in-sync with the measured arterial pressure. The upstream occlusion is induced at time approximately t = 7 seconds, resulting in a decay in pressure as blood drains from the arterial tree. The pressure oscillations, associated with myocardial contraction, continue during the decay. The final pressure attained after the approximately four-second transient decay is denoted the zero-flow pressure, Pzf, which is used as a metric of myocardial resistance vessel tone. In our analysis we match model simulations to the full time course of the zero-flow pressure experiment, as illustrated in Figure 1.

**Figure 1.**
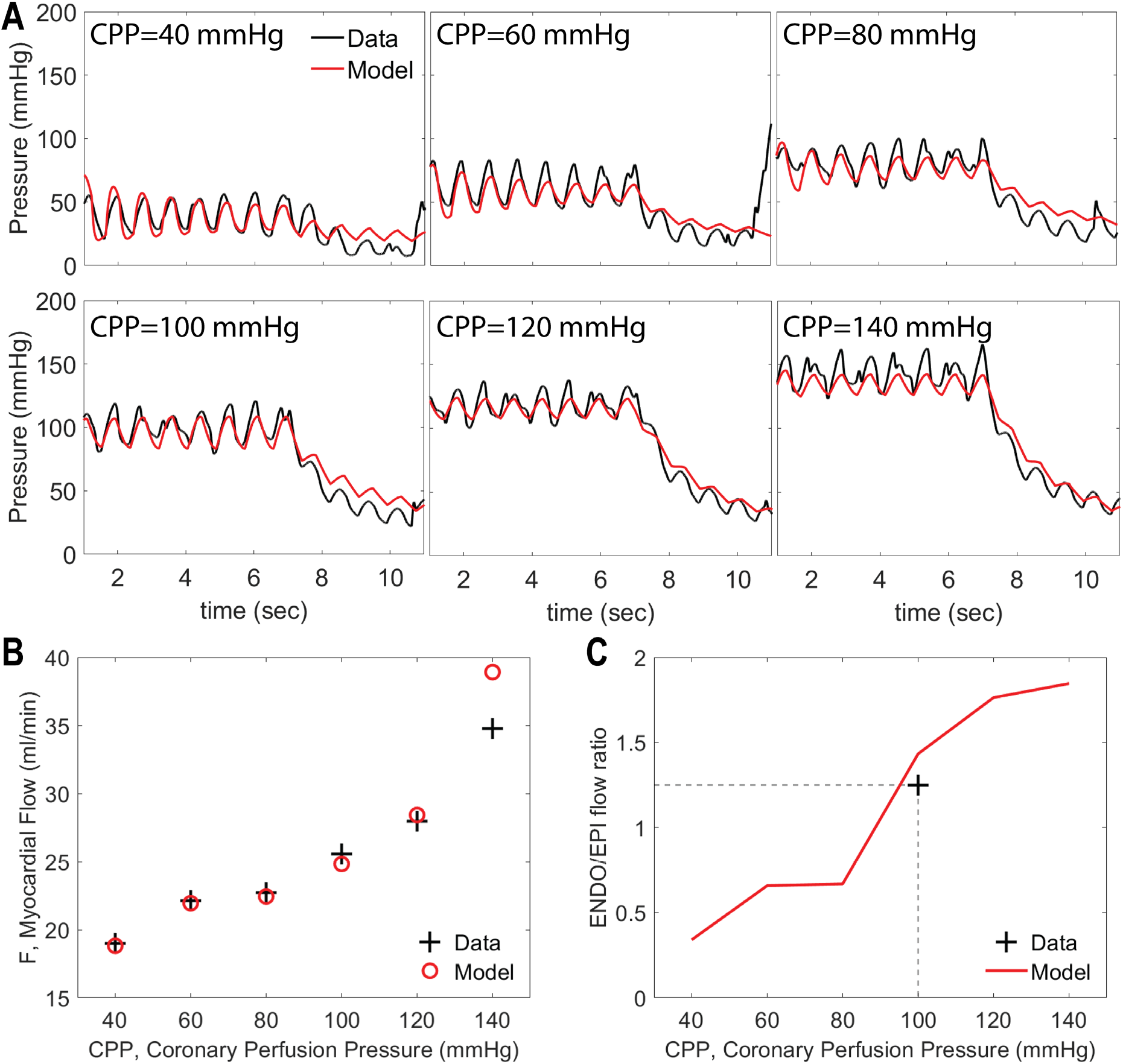
An example data set used to identify Model 1 for an individual animal under control conditions. A. Pressure recordings obtained at 6 levels of CPP following occlusion of left anterior descending (LAD) artery. The artery is occluded at ~7 sec and the decay distal to the occlusion is recorded. B. Total LAD flow, measured at different values of CPP. D. Myocardial subendocardial-to-subepicardial (ENDO/EPI) flow ratio before the occlusion. Data in Panels A and B are from Kiel et al. [4]. Measured ENDO/EPI flow ratio under baseline control conditions is targeted to 1.25 (see text for details). Measured data in panels is plotted in black; model simulations in red.

Experiments from Kiel et al. also provide data on total flow (before occlusion) at each level of initial perfusion pressure, as illustrated in Figure 1B for the baseline experimental condition for an individual animal. Finally, our model identification is constrained to match the experimentally measured baseline subendocardial-to-subepicardial flow ratio (ENDO/EPI), as illustrated in Figure 1B.

The zero-flow pressure experiment was repeated under three different experimental conditions: (1) control; (2) hemodilution; and (3) hemodilution + dobutamine. Hemodilution was induced by gradually replacing equal volumes of blood with a synthetic plasma expander. Dobutamine was administered by an intravenous drip to increase heart rate to ~75–100% above baseline levels.

Additional experimental data on conscious animals are used for validation. Briefly, animals were chronically instrumented with an arterial catheter, a perivascular flow transducer around the left anterior descending coronary artery, and with a solid state pressure transducer in the left ventricle as previously described [8]. Following recovery from surgery, hemodynamic variables were continuously measured under baseline resting conditions and subsequently during treadmill exercise.

### Overview of Model Formulation

Coronary blood flow regulation is simulated by integrating a lumped model for myocardial perfusion in subendocardial, midwall, and subepicardial layers of the myocardium (Model 1) with a model of resistance vessel mechanics accounting for myogenic, autonomic, and metabolic regulatory mechanisms (Model 2). In practice, the integrated model is identified in a two-step process. First, the lumped model is fit to data from zero-flow pressure experiments, yielding estimates of vessel diameters as functions of transmural pressure in the different myocardial layers under different levels of perfusion pressure and oxygen demand, at different heart rates, and under different experimental conditions. Next, the vessel mechanics model (Model 2) identified by comparing a panel of model formulations representing competing hypotheses for the metabolic mechanism to estimated diameters obtained from Model 1. Finally, the identified vessel mechanics model is validated by combining the two models to simulate myocardial blood flow in exercise versus baseline resting conditions.

### Three-Layer Lumped Model of Myocardial Perfusion (Model 1)

To simulate myocardial perfusion, we adapt the lumped model developed by Spaan et al. [9] and Mynard et al. [5, 10], illustrated in Figure 2. For the zero-flow pressure experiments, this model is used to simulate constant-pressure blood flow to the myocardium driven to the LAD by the extracorporeal servo-controlled pump, and the zero-flow pressure decay transient. The hydraulic properties of the perfusion tubing and epicardial vessels are governed by a series lumped resistance, blood inertance, and compliance, as illustrated in Figure 2. A venous resistance and compliance drain into the right-atrial pressure (P_RA_) downstream of a valve that prevents backflow in low perfusion pressures [11]. The myocardial circulation is divided to three parallel circuits representing subepicardial, midwall, and subendocardial layers. Within each layer, three serial resistances represent arterial, capillary, and venous compartments. Using Poiseuille resistance-volume relationship, the arterial and venous resistances (*R*_*1*_ and *R*_*2*_ in Figure 2) are computed

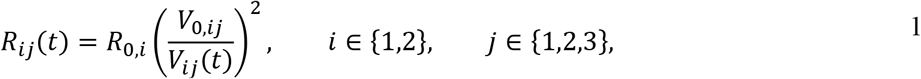

where *R*_0,*i*_ and V_0,*i*_ are reference resistance and volume, respectively, and the index j denotes myocardial layer: (1. subepicardial, 2. midwall, and 3. subendocardial). The instantaneous volume *V*_*ij*_ is (volume of compartment *i* in layer *j*) computed as

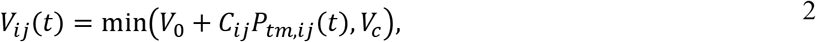

where transmural pressure (*P*_*tm,ij*_) is the difference between to blood pressure (*P*) and the intramyocardial pressure (*P*_*im,j*_) [5]. Based on the analysis in [5], we assume

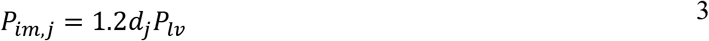

where *d*_*j*_ is the relative depth of subendocardial (*d*_*j*_ = 5/6), midwall (*d*_*j*_ = 1/2) and subepicardial layers (*d*_*j*_ = 1/6). To estimate *P*_*lv*_, we use half-sine functions constrained to match the measured aortic pressure where *P*_*lv*_ = *p*_*ao*_ in the systolic phase and *P*_*lv*_ = 5mmHg in diastolic phase. An example estimated ventricular pressure time course and corresponding measured aortic pressure time course are illustrated in Supplementary Figure S1.

**Figure 2.**
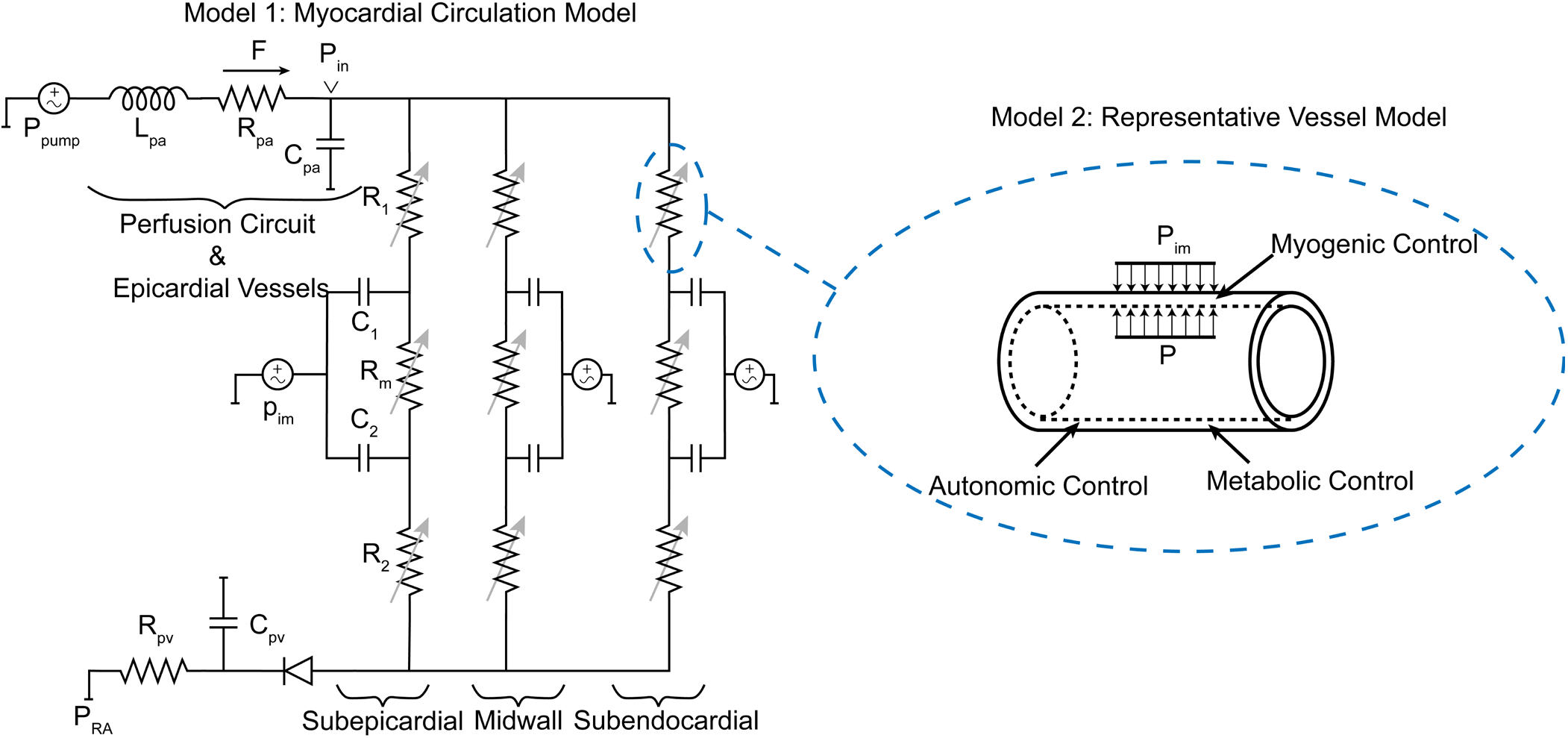
Schematic of the modeling approaches. Model 1: The myocardial circulation model which determines the flow and pressure in each layer of the myocardium. Model 2: A representative vessel model endowed with regulatory mechanisms to determine the level of vascular tone in each layer.

This model formulation allows for a potential collapse of vessels in cases of largely negative transmural pressures. This collapse is prevalent in the subendocardial layer and is characterized with a transmural pressure below which the vascular volume remains almost constant (*V*_*c*_) [4, 12].

Lastly, the middle resistance *R*_*mj*_ in each layer is computed using the resistance of the arterial and venous compartments

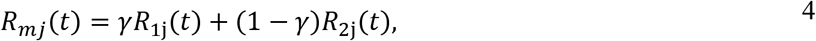

where *γ* is a parameter between 0 and 1. To ensure a physiologically realistic transmural flow, the reference resistance *R*_0,1_ and the arterial compliances *C*_1*j*_ are scaled with two factors: subepicardial to subendocardial resistance factor *r*_*f*_ and compliance factor *c*_*f*_.

To simulate changes in vascular tone (dilation/constriction) in response to changes in CPP in the lumped model a factor is *f*_*CPP*_ is used such that

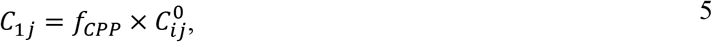

where *f*_*i*_ > 1 represent vasodilation for CPP<100, *f*_*j*_ < 1 vasoconstriction for CPP>100, and 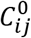 is a reference value corresponding to the baseline conditions taken to be the CPP = 100 mmHg. The vasoreactivity factor is assumed to vary across different across the myocardium. To incorporate graded vasoreactivity, a parameter 0 < *β* < 1 is introduced such that

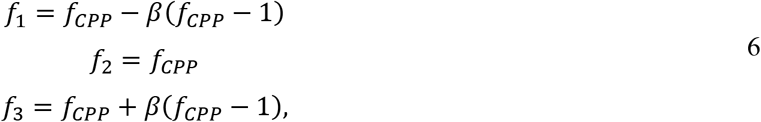

where *f*_*CPP*_ is the overall vasoreactivity factor.

To simulate the zero-flow pressure experiment, we divide the experiment to two parts: part 1 when the pump is driving flow to the LAD perfusion region; and part 2 when the pump is off and the extracorporeal circuit is clamped. To simulate first part of the experiment, we assume that the output pressure from the pump is constant and set to the nominal CPP for simulation time 0 < *t* < *t*_off_, where *t*_off_ when the pump is turned off and the extracorporeal circuit is clamped. For this first part of the experiment, the fixed driving pressure is used to drive the model and the resulting simulated flow is compared to the measured data.

To simulate the second part of the experiment, flow is imposed, and the resulting pressure transient is simulated. Specifically, since the model cannot accommodate a step change in flow, a rapid continuous decrease to zero flow is imposed using the function

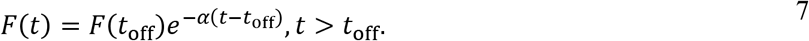

The rate of decay is set to the value *α*= 10 sec^−1^, to obtain a decay to zero flow that is effectively instantaneous compared to the timescale of the observed pressure decay.

#### Parameter Estimation for Model 1

Phasic tracings of perfusion pressure and flow for each experimental preparation are used to calibrate the myocardial circulation model. The perfusion pressure is measured continuously in both steps of the experiment using a cannula in the left anterior descending (LAD) coronary artery in both steps of the experiment. Accordingly, in our model the pressure measurements correspond to *P*_*in*_ in Figure 2. The model residual vector is defined as

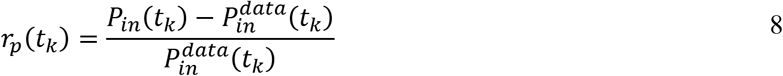

where 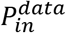 is the measured data at time *t*_*k*_. The flow across *R*_*pa*_, *q*_*in*_, is compared to the measured flow. Due to pump-induced noise in flow tracings, the average flow before closing the circuit 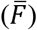 is used in the objective:

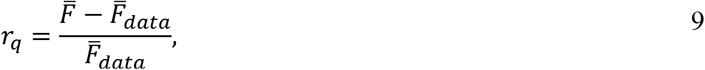

where 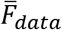 the average of flow before the extracorporeal circuit is clamped. The cost function for the parameter estimation is a weighted sum of the error in time-resolved pressure tracings, average flow before clamping the perfusion circuit, and a penalty term to ensure a physiological subendocardial to subepicardial flow ratio (ENDO/EPI) for each CPP level

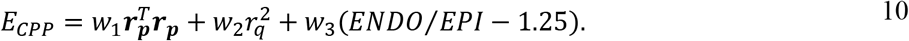

The weights *w*_1_ and *w*_2_ are chosen such that the contribution of the pressure and flow to the cost function are the same order. The weight *w*_3_ is varied for CPP levels, assuming the largest weight for the CPP = 100 mmHg case, where the target ENDO/EPI ratio at rest is 1.25, reflecting the approximate average of values reported in the literature [13–15]. Finally, the total cost function for parameter estimation can be written as

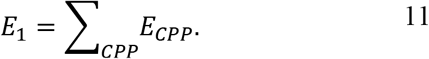

In this work, the model parameters corresponding to the venous components of the model were fixed, and the arterial component parameters were adjusted to minimize the objective function *E*_1_. Table 1 lists the adjustable parameters and their description.

**Table 1.** List of adjusted parameters for Model 1.

| Parameter | Mean and (Range) of Estimated Values | Unit | Description |
| --- | --- | --- | --- |
| $L_{pa}$ | 1.737 (0.129-3.976) | mmHg / (mL s<sup>-2</sup>) | Blood inertance in perfusion circuit and epicardial vessels |
| $R_{pa}$ | 2.729 (0.367-8.511) | mmHg / (mL s<sup>-1</sup>) | Resistance in perfusion circuit and epicardial vessels |
| $C_{pa}$ | 0.0211 (0.009-0.042) | mL / mmHg | Compliance in perfusion circuit and epicardial vessels |
| $V_0$ | 0.455 (0.218-0.643) | mL | Reference volume |
| $V_c$ | 0.194 (0.020-0.599) | mL | Collapsible volume |
| $R_{0,3}$ | 104.00 (46.33-217.43) | mmHg / (mL s<sup>-1</sup>) | Reference subendocardial arterial resistance |
| $C_{13}$ | 0.0044 (0.001-0.010) | mL / mmHg | Subendocardial arterial compliance |
| $c_f$ | 0.8686 (0.716-0.999) | - | The ratio of subepicardial to subendocardial compliance |
| $r_f$ | 1.4549 (2.047-1.006) | - | The ratio of subepicardial to subendocardial resistance |
| $\beta$ | 0.3360 (0.034-0.819) | - | Degree of graded vasoreactivity |
| $f_{CPP}$ | [7.43 (1.006-19.522),<br>7.11 (2.015-12.183),<br>2.55 (1.163-6.221),<br>0.38 (0.002-0.999),<br>0.30 (0.003-0.999)] | - | Vasoreactivity factor for CPP=40,60,80,120,140<br>mmHg |

### Representative Vessel Model (Model 2)

In the representative vessel model, the arterial component in each layer is represented by a single vessel endowed with passive and active tension models as well as myogenic, metabolic, and autonomic regulatory mechanisms. Following [6, 16], the vessel wall tension is sum of a passive tension *T*_*pass*_ and an active tension *T*_*act*_, which is the tension generated by the vascular smooth muscle cells (VSMs). The active tension is determined as the product of the maximal active tension 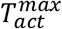 in a given vessel and the activation level *A*. Thus, the vascular tension can be written as

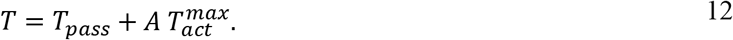

The passive response of the vessel is expressed as a nonlinear function of diameter, by rearranging the nonlinear pressure-diameter relation in [17]

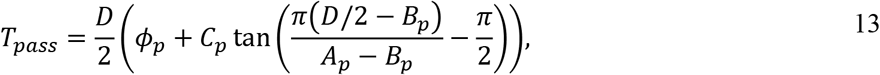

where *A*_*p*_ and *B*_*p*_ represent the asymptomatic maximum and minimum radii, *C*_*p*_ is the bandwidth between *A*_*p*_ and *B*_*p*_, and *ϕ*_*p*_ corresponds to the pressure offset determining the half-way point between *A*_*p*_ and *B*_*p*_. The maximal active tension generated by VSM is expressed by modifying the Gaussian function given in [6] so that the maximal possible active tension diminishes for small diameters.

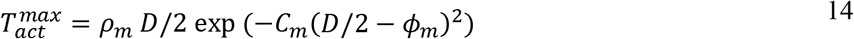

Following [16], the activation level *A* depends on the total stimulus *S*_*tone*_ in a sigmoidal manner

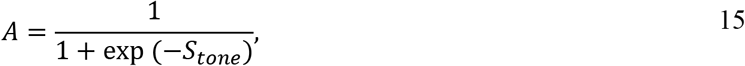

where *S*_*tone*_ is a net summation of stimuli from the myogenic *S*_*myo*_, metabolic *S*_*meta*_, autonomic *S*_*HR*_ mechanisms, and an offset *C*_0_:

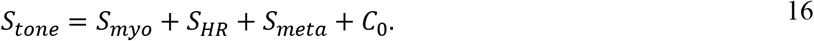

We assume myogenic response is a linear function of the vascular tension, so we can write

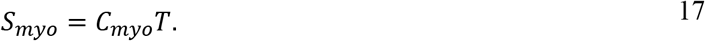

The autonomic response is assumed to be a function of the heart rate and to have only dilatory effects on the myocardial vessels. Moreover, a baseline offset *HR*_0_ is used to determine the threshold for autonomic signal generation.

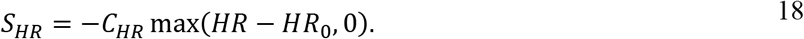

Similarly for metabolic mechanism, we assume the stimuli linearly depends on the signal

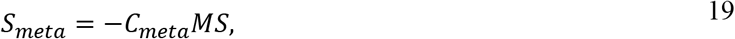

where *MS* is the metabolic signal, and the negative sign corresponds to a vasodilatory response.

In this work we evaluated seven different representations of the metabolic signal:

1. *ATP-dependent (ATP)*: ATP release from red blood cells provides the metabolic feedback signal for local control of perfusion. The ATP transport model of Pradhan et al. [18] accounts for oxygen saturation-dependent leak of ATP from red blood cells and a flow-dependent washout of ATP from the circulation. From Pradhan et al., the metabolic signal is assumed proportional to venous ATP, which is governed by

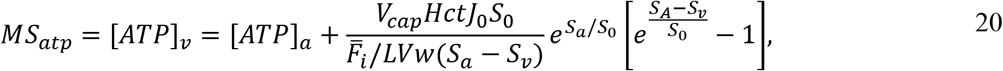

where [*ATP*]_*v*_ and [*ATP*]_*a*_ are arterial and venous plasma ATP concentrations, *V*_*cap*_ is the capillary volume density, *LVw* is the left ventricle weight, *J*_0_ is the ATP release rate in coronary vascular bed, and *S*_0_ is the ATP release parameter of the model.
2. *Myocardial oxygen consumption (MVO*_*2*_): Myocardial oxygen consumption determines the metabolic signal. Under given experimental conditions, total MVO_2_ in is approximated

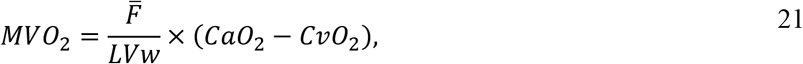

where *CaO*_2_ and *CvO*_2_ are the arterial and venous oxygen content. Experimental studies have shown that that myocardial fiber shortening and energy demands in subendocardium are higher than those of subepicardium [19]. This leads to variable oxygen consumption levels across the myocardium, with subendocardial to subepicardial MVO_2_ ratio (*R*_*M*_) of ~1.5 [19, 20]. Based on these observations, the layer-wise MVO_2_ is computed by dividing MVO_2_

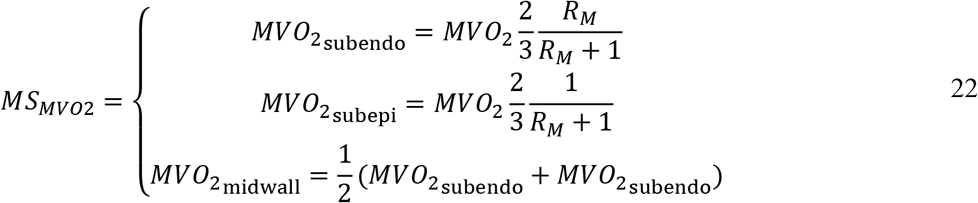
3. *Oxygen extraction (S*_*a*_ − *S*_*v*_): The arterial-venous difference in oxygen saturation acts as a signal for metabolic control.

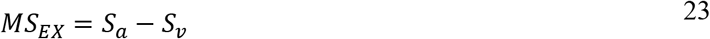
4. *Layer-wise variable oxygen extraction (LS*_*v*_): The regional arterial-venous difference in oxygen saturation acts as a signal for metabolic control. Using the layer-wise MVO_2_, we compute venous oxygen saturation and arterial-venous oxygen extraction in each of three layers of the myocardium.

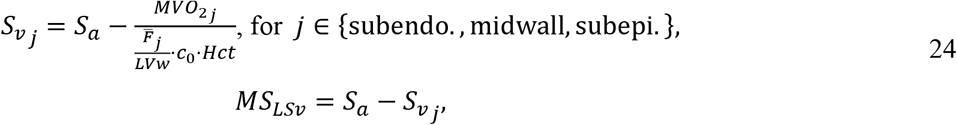

where *c*_0_ is the oxygen content of fully oxygen saturated oxyhemoglobin in a red blood cell.
5. *Flow times oxygen extraction (FΔS)*: The metabolic signal is the product flow and oxygen extraction in each layer of the myocardium.

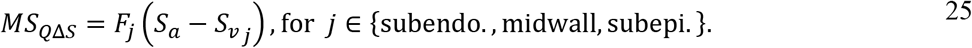
6. *Flow times MVO*_*2*_ *(FM)*: The metabolic signal is the product flow and MVO_2_ in each layer of the myocardium.

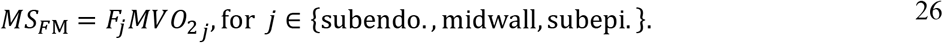
7. *Nonlinear metabolic rate model (MVO*_*2*_^*2*^): The metabolic signal is the square of the myocardial oxygen consumption rate.

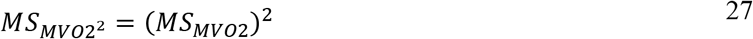

#### Parameter Estimation for Model 2

Outputs of the parameterized Model 1, fit to data from the zero-flow pressure experiments, are used to estimate the layer-wise average diameters of representative resistance vessels, 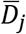, for each experimental condition (control, hemodilution, hemodilution+dobutamine), each level of CPP, in each layer of the myocardium, and for each of four experimental animals. Additional inputs to Model 2 are average hemodynamics (pressure and flow) and estimated tissue pressures in each layer of the myocardium, blood oxygenation measurements, and heart rate.

The layer-wise average flow 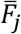, transmural pressure *P*_*tm,j*_, and resistance 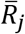 are computed from the cycle averaged simulated of the lumped Model 1 under each experimental condition for each experimental animal. The estimated model-1 resistances 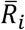 of the arterial components are used to determine representative vessel diameters:

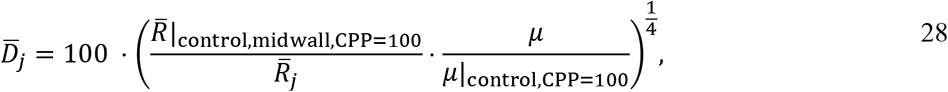

where the diameter of midwall at CPP=100 mmHg level is assumed to be 100 μm. The effective blood viscosity *μ* determined as a function of hematocrit using the relation [21]:

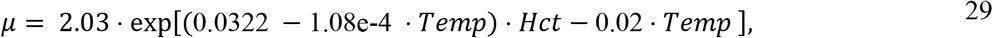

where *Hct* and *Temp* are hematocrit and temperature, respectively.

Figure 3A shows how the estimated representative vessel diameter varies with transmural pressure in the midwall layer of an individual animal for the three different experimental conditions. To match the predictions of the vessel model (Model 2) to these estimated diameters, the following objective function is used for parameter estimation

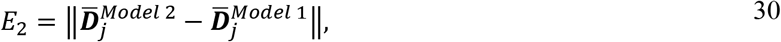

where 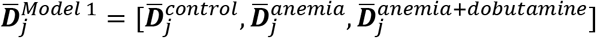 is the vector of computed diameters representing the same pig/same layer, and 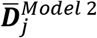 is the corresponding Model 2 predictions. The adjustable parameters are listed in Table 2.

**Figure 3.**
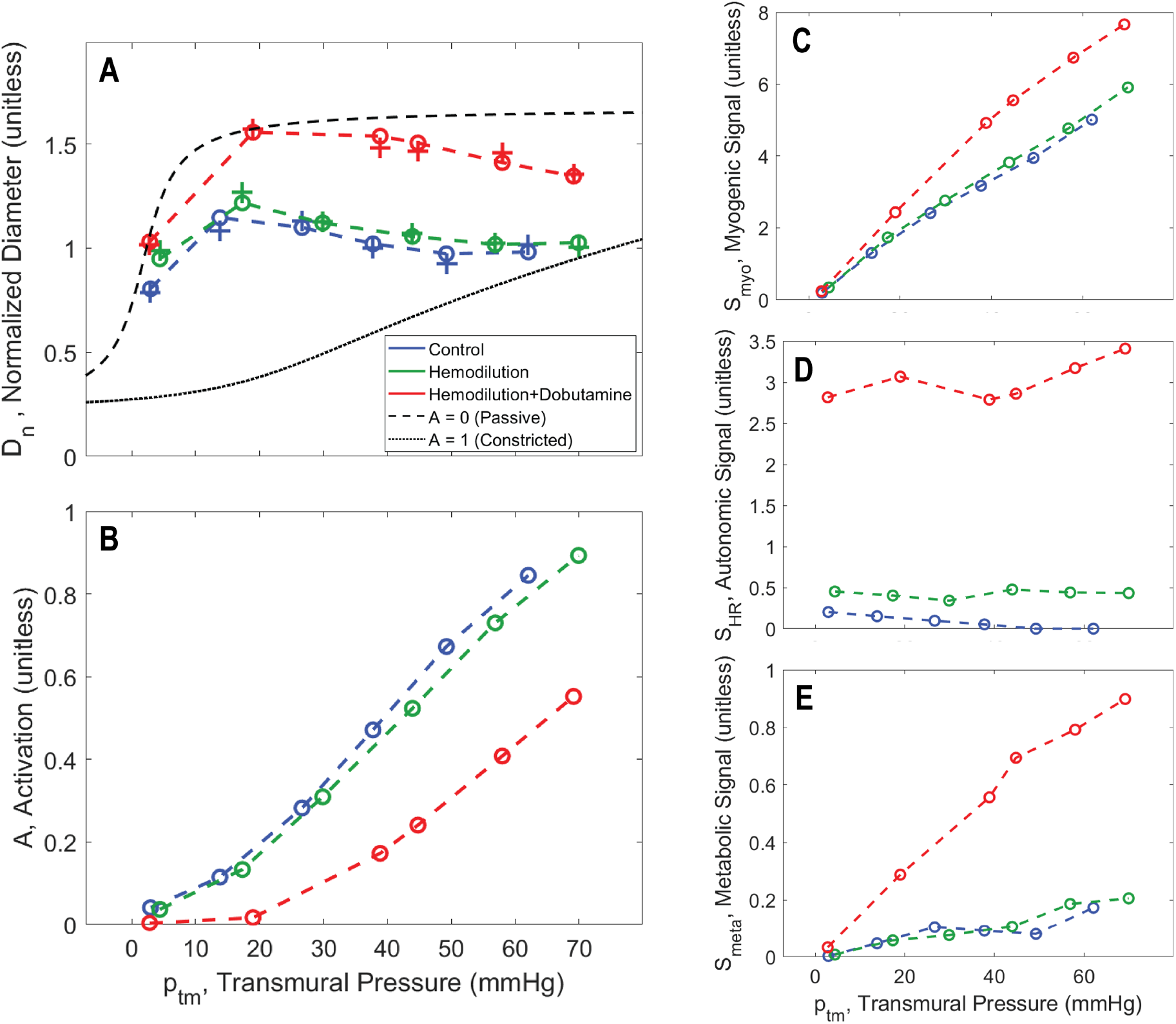
Vasoregulation as a function of transmural wall pressure. A. Model predictions of relative resistance vessel diameters are shown for the midwall layer for pig C for the midwall layer. Diameters for the three experimental conditions, associated with the fits of Model 1 to the zero-flow pressure experiment, are plotted as “+” markers: blue for control; green for hemodilution; and red for hemodilution + dobutamine. The matches of Model 2-based predictions (using the ‘*F·M’*, flow times MVO_2_, metabolic signal) to the diameter estimates are plotted as “o” markers connected by dashed lines. B. Predicted total smooth muscle activation is plotted as a function of transmural pressure in the zero-flow pressure experiment. C. Predicted myogenic activation signal is plotted as a function of transmural pressure in the zero-flow pressure experiment. D. Predicted autonomic activation signal is plotted as a function of transmural pressure in the zero-flow pressure experiment. E. Predicted metabolic activation signal is plotted as a function of transmural pressure in the zero-flow pressure experiment.

**Table 2.**
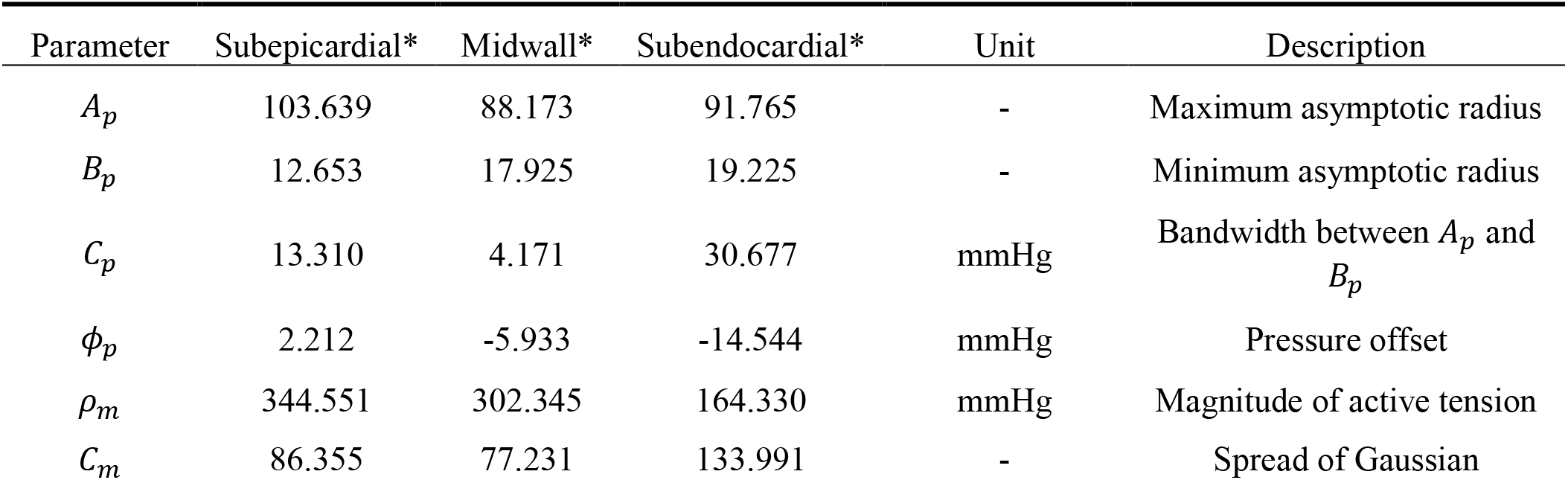
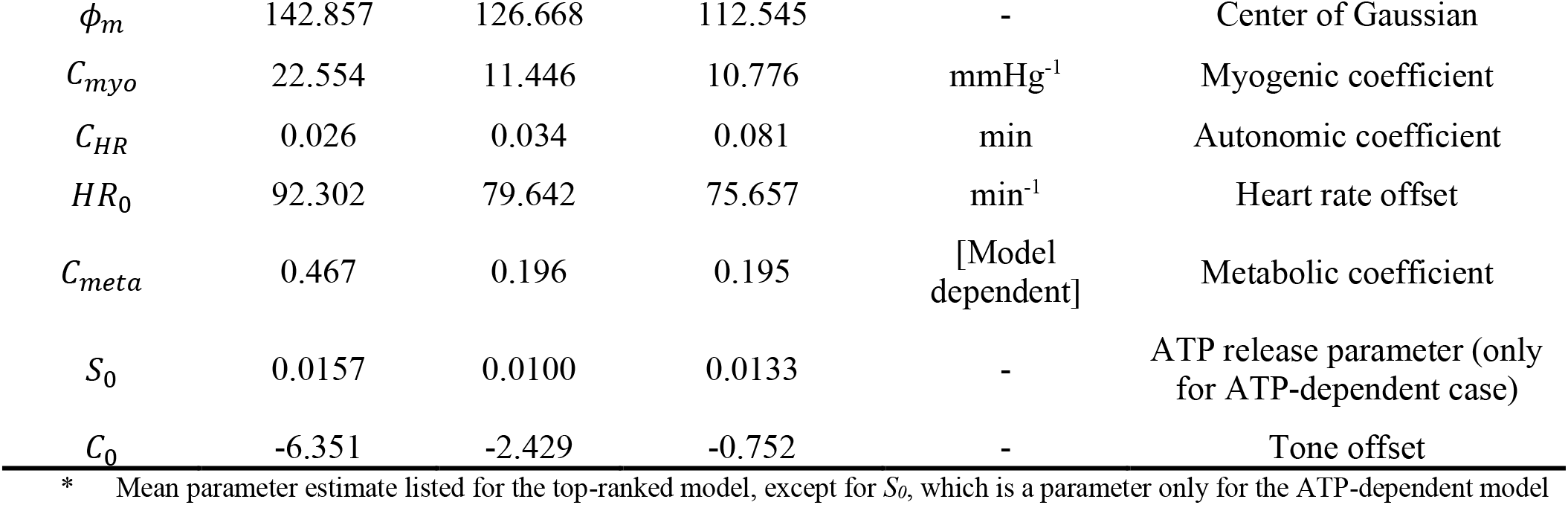
List of adjustable parameters for Model 2

| Parameter | Subepicardial* | Midwall* | Subendocardial* | Unit | Description |
| --- | --- | --- | --- | --- | --- |
| $A_p$ | 103.639 | 88.173 | 91.765 | - | Maximum asymptotic radius |
| $B_p$ | 12.653 | 17.925 | 19.225 | - | Minimum asymptotic radius |
| $C_p$ | 13.310 | 4.171 | 30.677 | mmHg | Bandwidth between $A_p$ and $B_p$ |
| $\phi_p$ | 2.212 | -5.933 | -14.544 | mmHg | Pressure offset |
| $\rho_m$ | 344.551 | 302.345 | 164.330 | mmHg | Magnitude of active tension |
| $C_m$ | 86.355 | 77.231 | 133.991 | - | Spread of Gaussian |
| $\phi_m$ | 142.857 | 126.668 | 112.545 | - | Center of Gaussian |
| $C_{myo}$ | 22.554 | 11.446 | 10.776 | mmHg<sup>-1</sup> | Myogenic coefficient |
| $C_{HR}$ | 0.026 | 0.034 | 0.081 | min | Autonomic coefficient |
| $HR_0$ | 92.302 | 79.642 | 75.657 | min<sup>-1</sup> | Heart rate offset |
| $C_{meta}$ | 0.467 | 0.196 | 0.195 | [Model<br>dependent] | Metabolic coefficient |
| $S_0$ | 0.0157 | 0.0100 | 0.0133 | - | ATP release parameter (only<br>for ATP-dependent case) |
| $C_0$ | -6.351 | -2.429 | -0.752 | - | Tone offset |

\* Mean parameter estimate listed for the top-ranked model, except for $S_0$ , which is a parameter only for the ATP-dependent model

#### Model 2 model selection

The parameter estimation is repeated using each of the seven different metabolic signal models. To compare the performance of each model formulation, we use a modified second order Akaike information criterion (*AICc*), which is used for model selection across multiple datasets with small sample size [22, 23]

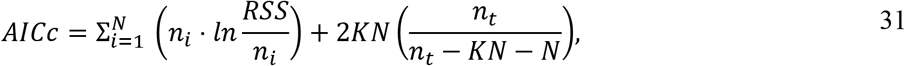

where *RSS* is the residual sum of squares 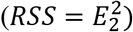, *K* is the number of parameters, *N* is the number of data sets (4 pigs × 3 layers = 12 datasets), and *n*_*i*_ is the number of data points in each data set (Pig 1: 15, Pig 2: 12, Pig 3: 18, Pig 4: 18) and *n*_*t*_ is the total number of data points (63). Then, a parameter Δ for each is computed as

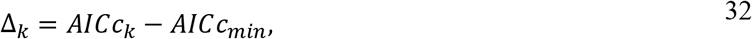

where *AICc*_*k*_ is the Akaike criterion for *k*^th^ model, and *AICc*_min_ is the minimum *AICc* value. Finally, the relative likelihood (*RL*_*i*_), is given by

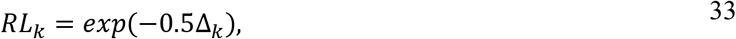

which is the probability that a particular *k*^th^ model is a better model than the model with the minimum *AICc* value.

### Simulating Coronary Flow in Rest versus Exercise Conditions

Once the coronary flow regulation model is constructed, the identified model framework, with Models 1 and 2 integrated together, is validated by simulating in vivo myocardial perfusion in resting and exercise conditions. These simulations are compared to data collected in conscious animals, for which aortic and left ventricular pressure and LAD flow were simultaneously measured. The measured aortic pressure was used to drive the model, and simulations of LAD flow compared to data. Since the parametric identification from the zero-flow pressure experiments yields different optimal parameter values for each individual animal, and since the conscious exercise experiments were conducted on different animals and under different experimental conditions, a subset of parameters was identified to adjust to match the conscious state data. The model formulation is validated by showing that the model formulation using the top-ranked metabolic mechanism is able to match the conscious state data markedly better than lower ranked models.

To simulate myocardial perfusion in vivo the two model components are simulated iteratively until they reach convergence. The measured aortic and left ventricular pressure data are inputs to Model 1 for a given set values of resistance vessel parameter values, *C*_11_, *C*_12_, and *C*_13_, representing vessel tones in the three myocardial layers. The predicted flow from Model 1, along with the given measured heart rate and estimated MVO_2_, is used to compute the metabolic signal for use with Model 2. Model 2 is simulated to estimate the vessel diameters in each layer, from which the average resistances and compliances are computed:

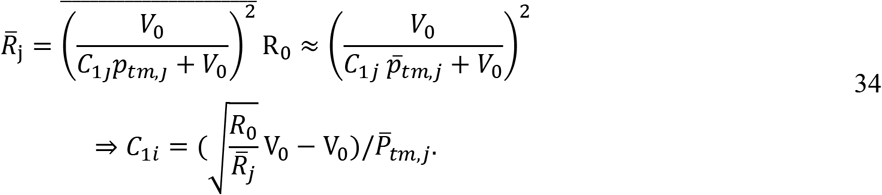

The computed compliance values are fed back into Model 1, and the iterative process is carried out until Models 1 and 2 agree and the predicted hemodynamic state reaches convergence.

To calibrate the model to simulate exercise, we perform the following analysis to identify a set of adjustable parameters that satisfy two criteria: (1.) parameters in the set are estimated from the zero-flow pressure experiments with a relatively high degree of uncertainty; (2.) simulations of in vivo myocardial perfusion are sensitive to the values of parameters in the set of identified parameters.

The uncertainty of the values estimated from the zero-flow pressure experiments is quantified based on the coefficient of variation *CoV*

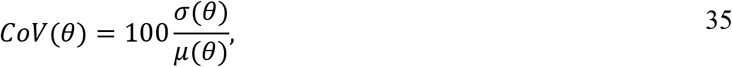

where *σ* and *μ* are the standard deviation and average of parameter *θ* used in Model 2. Next, we calculate parameter sensitivities using rest and exercise simulations [18]

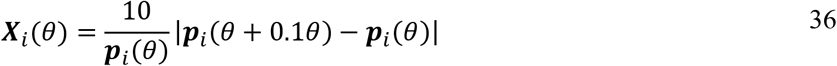

where prediction vectors ***p***_*i*_ are simulation results; ***p***_1_ = *F*_*rest*_(***t***), ***p***_2_ = *F*_*exer*_.(***t***), ***p***_3_ = (*ENDO/EPI*)_*rest*_, ***p***_4_ = (*ENDO/EPI*)_*exer*_. correspond to flow and ENDO/EPI flow ratio in rest and exercise. Finally, the model sensitivity *S* to each parameter is computed is the sensitivity of each prediction

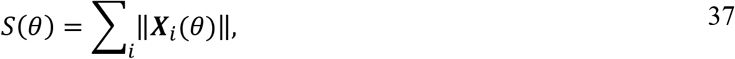

To find the set of adjustable parameters, we find the parameters *θ* with *CoV*(*θ*) > 30% and *S*(*θ*) > 0.1. To estimate the parameters, we define model residual errors as

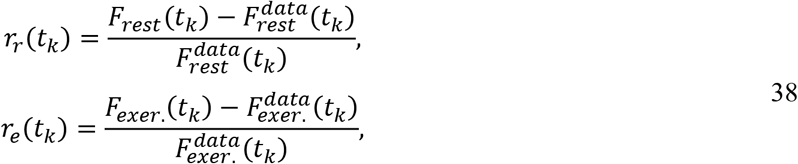

where 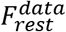 and 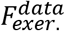 are LAD flow data in rest and exercise. Then, the cost function for the parameter estimation is

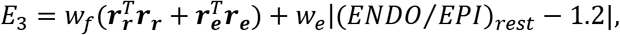

with weights *w*_*f*_ and *w*_*e*_.

## Results

### Model 1 Identification

Figure 1 shows an example data set for the zero-flow pressure experiment along with a fit of model 1 for an individual animal under control conditions. The top two rows show the pressure tracings at different levels of CPP. At roughly t = 7 seconds, the LAD flow is occluded and the decay of pressure at a distal site is recorded for approximately 4 seconds. The bottom panels show data and model predictions of average flow and ENDO/EPI flow ratio measured before the occlusion. Supplementary Figures S2-S13 show model fits to the zero-flow pressure time course data for all animals and experimental conditions.

Figure 4 shows the measured and model-predicted total flow data for the four individual pigs for the three different experimental preparations. In control and hemodilution conditions flow is maintained at roughly constant values over an autoregulatory range of CPP values between 60 and 120 mmHg. The total flow is elevated in hemodilution compared to control, in order to maintain oxygen delivery. Infusion of dobutamine significantly increases the flow compared to the control and hemodilution conditions and tends to abolish the autoregulatory response in all animals.

**Figure 4.**
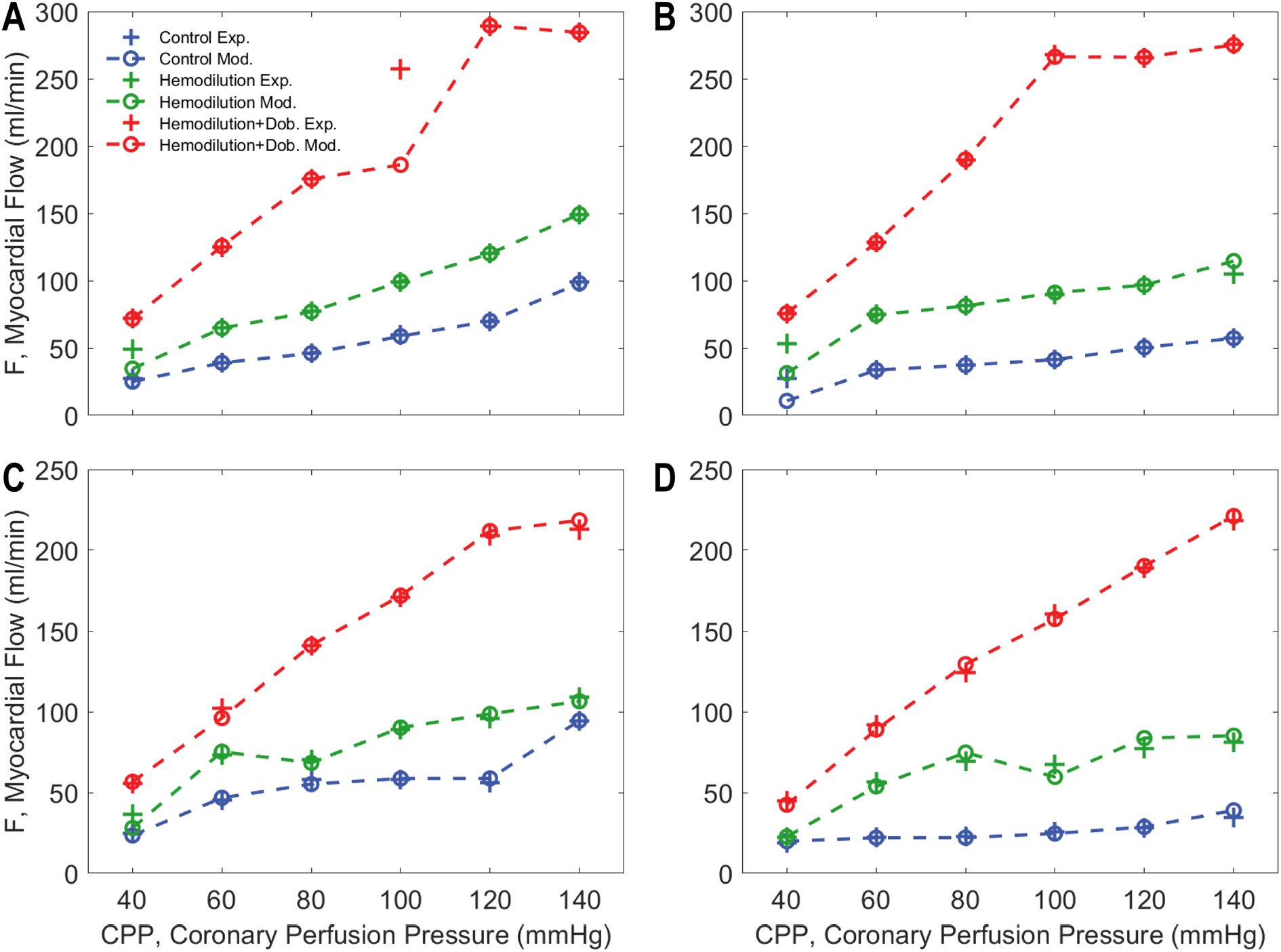
Measured versus model-predicted LAD flow for each of four experimental animals under three different experimental conditions. In control and hemodilution conditions, coronary autoregulation impedes a significant increase in flow with increases in CPP (60-120 mmHg). However, the overall myocardial blood flow is elevated with hemodilution. Dobutamine infusion significantly increases the flow compared to the control and hemodilution conditions and largely abolishes the autoregulatory response in all animals.

The Model-1 fits yield estimates of resistance vessel diameter at each CPP level under each experimental condition for each animal. These diameter estimates are illustrated in Figure 3 for all three experimental conditions for the midwall layer for one experimental animal. Analogous predictions for all myocardial layers and for all animals are plotted in Supplementary Figures S14-S25.

### Model 2 (Representative Vessel Model) Identification

As illustrated in Figure 3A and in Supplementary Figures S14-S25, Model 1 yields estimates of resistance vessel diameter as functions of transmural vessel pressure for three different experimental conditions, for three myocardial layers, and four individual animals at each of six values of CPP. Each of the seven competing versions of Model 2 (representing different formulations of the metabolic mechanism) are fit to this data set to determine the most likely model best able to represent the data. Example fits, associated with the flow-times-MVO_2_ metabolic mechanism, are shown in Figure 3A.

Table 3 lists the estimated AICc_k_, Δ_k_, and RL_k_ values for the seven models, illustrating that the flow-times-MVO_2_ metabolic mechanism is the highest ranked model, identified as the maximally likely among the models tested.

**Table 3.**
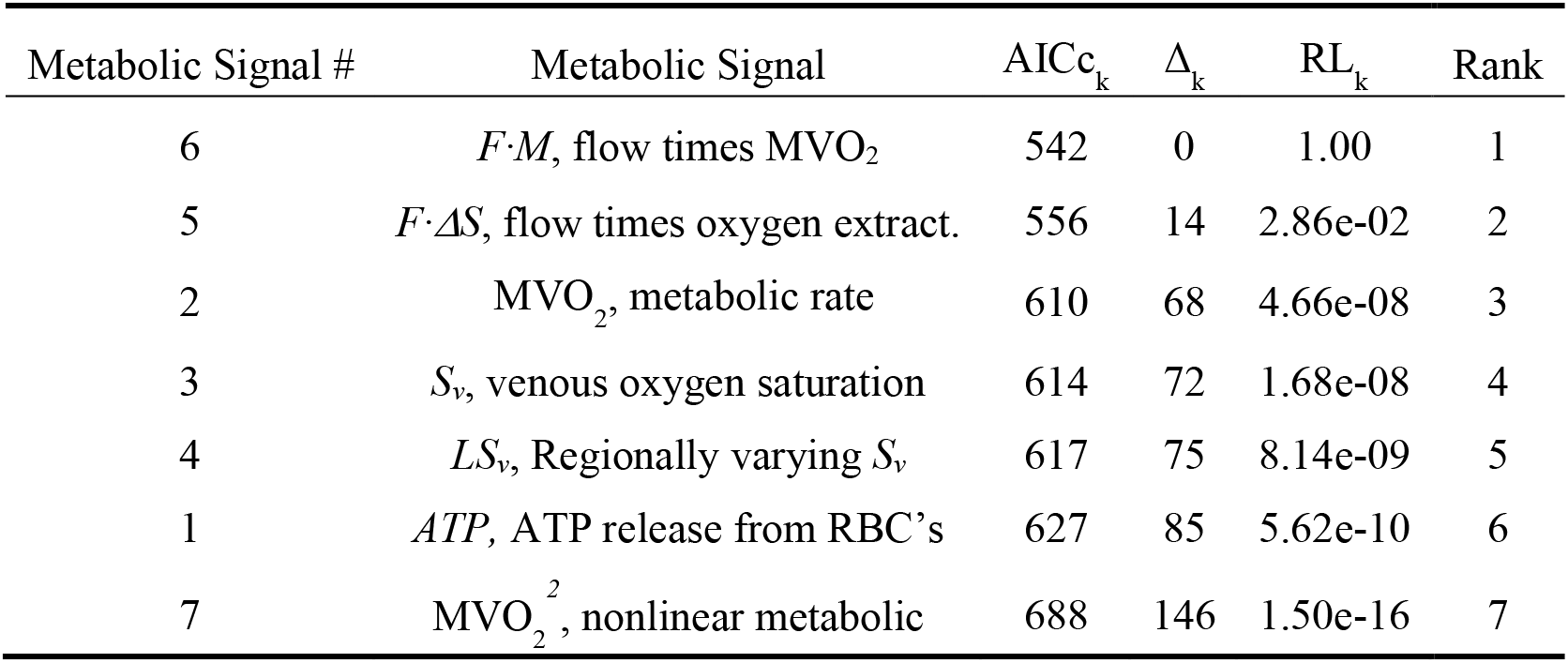
Models are ordered by their *AICc* value with the best model ranked 1.

Figure 3 also plots the regulatory signals predicted by the model for the fits to the midwall myocardial layer show in panel A. For all three experimental conditions, the resistance vessel constricts with increasing the transmural pressure. In control and hemodilution conditions, the total activation level *A* steadily increases from 0 for low CPPs (full dilation) to 0.8-0.9 for high CPPs. With the addition of dobutamine, the overall activation is shifted down. Panels C-E show the variation of each mechanism in response to changes in CPP and experimental conditions. The myogenic contribution increases with increasing the transmural pressure, with the greatest values in hemodilution+dobutamine case. The predicted autonomic contribution is nearly constant over the observed range of CPP because the heart rate is nearly constant for each experimental condition. The autonomic contribution is elevated in the hemodilution+dobutamine case due to a higher the heart rate for this condition compared to other cases.

The metabolic signal is predicted to be an order of magnitude smaller than the myogenic and autonomic signals for the midwall layer for this animal under these conditions. Panel C shows that the strongest contributor to the pressure autoregulatory response is the myogenic mechanism, with S_myo_ increasing from 0 at the lowest CPP to 8 at the CPP = 140 mmHg (corresponding to average transmural pressure of approximately 60 mmHg in the midwall region). The autonomic signal makes the biggest contribution to increasing diameter under hemodilution+dobutamine compared to other cases. This prediction is similar to that of Pradhan et al. [18], who predicted that the open-loop autonomic signal contributes more to the response in coronary flow to increasing demand in exercise than the metabolic feedback signal.

Figures 5, 6, and 7 illustrate how the predicted resistance vessel diameters and regulatory signals vary with CPP in each layer of the myocardium, summarizing predictions for all experimental animals. In the subepicardium (Figure 5), diameter decreases with increasing CPP under all experimental conditions. This trend is reversed in the subendocardium (Figure 7), where an increase in vasodilation with increasing perfusion pressure counters the action of increasing transmural pressure. In the hemodilution+dobutamine experiments, in particular (Figure 7B), the subendocardium is predicted to be nearly fully vasodilated across all values of CPP.

**Figure 5.**
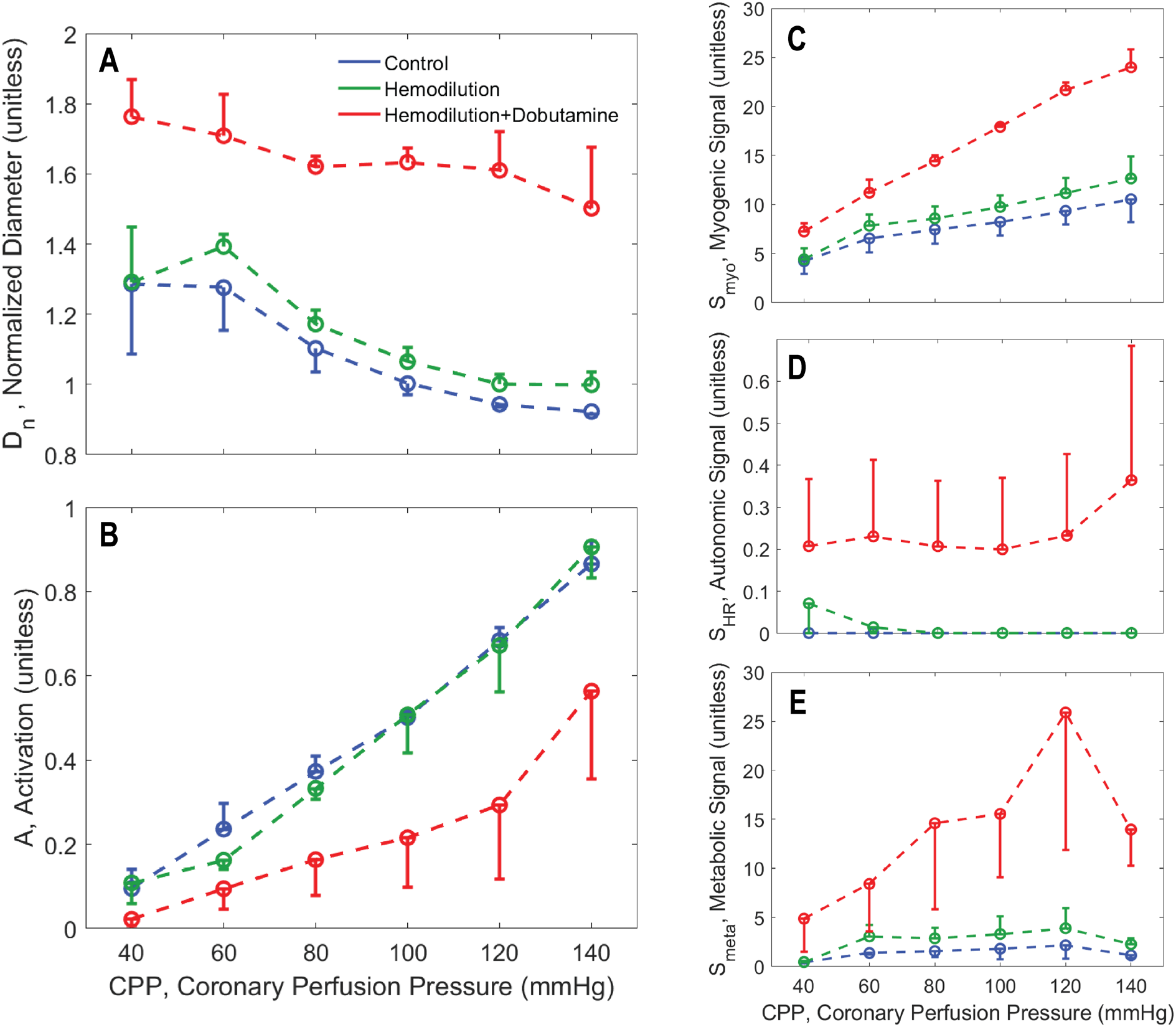
Vasoregulation in subepicardial vessels. A. Predicted mean and standard error of subepicardial resistance vessel diameter is plotted as a function of CPP. B. Predicted mean and standard error of total vessel activation is plotted as a function of CPP. C. Predicted mean and standard error of myogenic activation signal is plotted as a function of CPP. D. Predicted mean and standard error of autonomic activation signal is plotted as a function of CPP. E. Predicted metabolic activation signal is plotted as a function of CPP.

**Figure 6.**
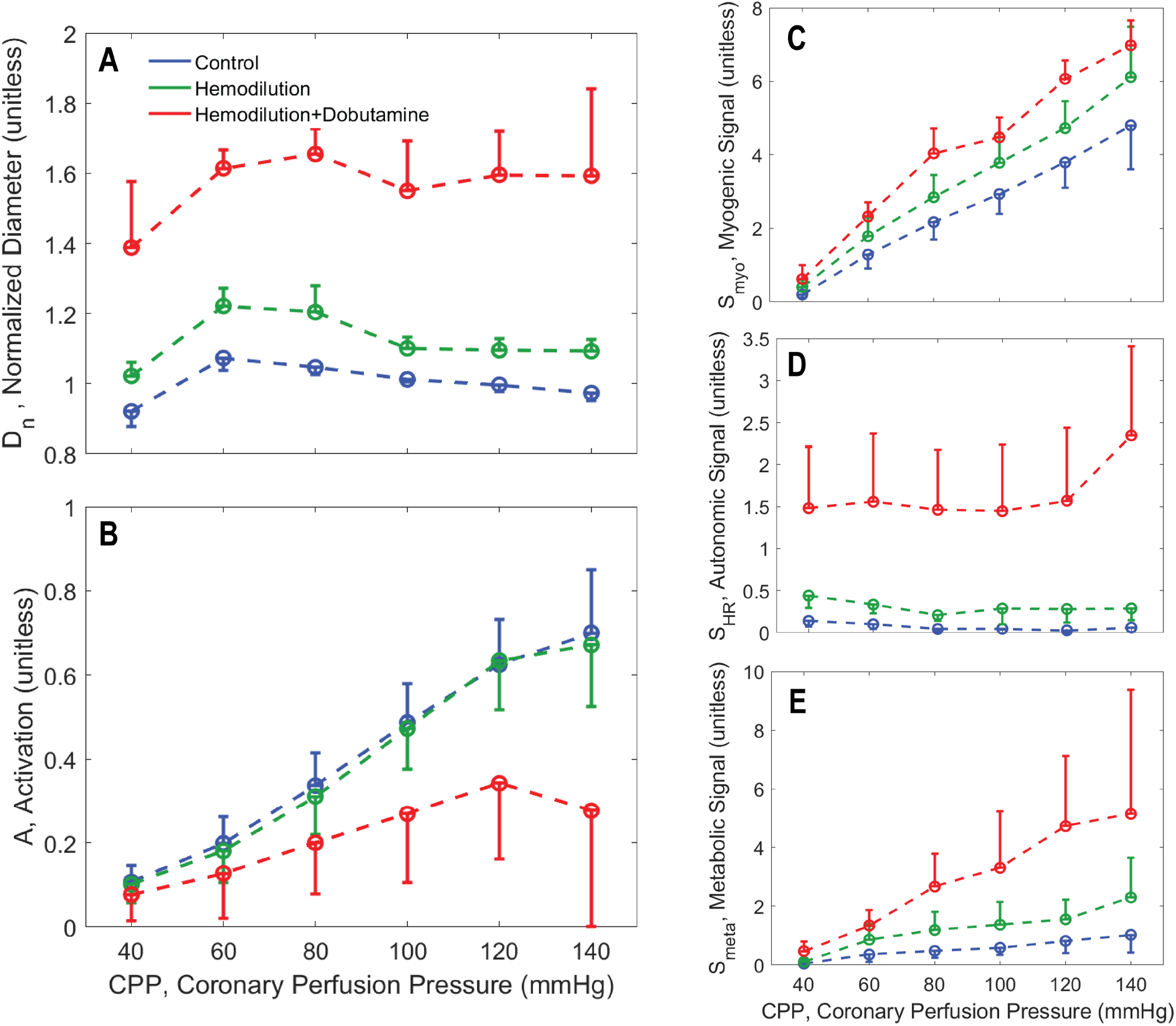
Vasoregulation in midwall vessels. A. Predicted mean and standard error of subepicardial resistance vessel diameter is plotted as a function of CPP. B. Predicted mean and standard error of total vessel activation is plotted as a function of CPP. C. Predicted mean and standard error of myogenic activation signal is plotted as a function of CPP. D. Predicted mean and standard error of autonomic activation signal is plotted as a function of CPP. E. Predicted metabolic activation signal is plotted as a function of CPP.

**Figure 7.**
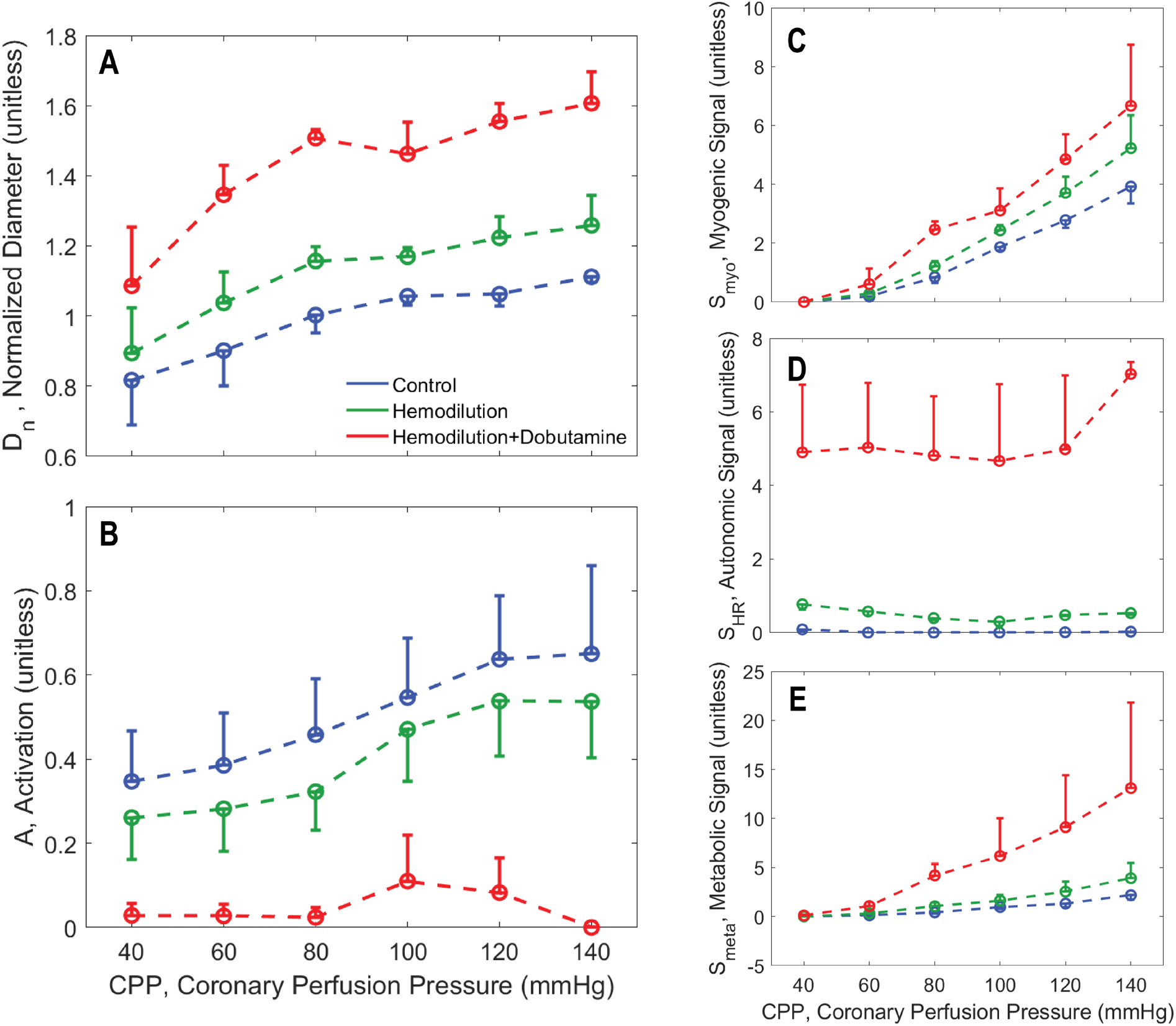
Vasoregulation in subendocardial vessels. A. Predicted mean and standard error of subepicardial resistance vessel diameter is plotted as a function of CPP. B. Predicted mean and standard error of total vessel activation is plotted as a function of CPP. C. Predicted mean and standard error of myogenic activation signal is plotted as a function of CPP. D. Predicted mean and standard error of autonomic activation signal is plotted as a function of CPP. E. Predicted metabolic activation signal is plotted as a function of CPP.

As expected, the myogenic mechanism (panel C of Figure 5, 6, and 7), makes the greatest contribution to the pressure autoregulatory response in the subepicardium, and smaller contributions in the midwall and subendocardial layers. The autonomic signal contribution (panel D of Figure 5, 6, and 7) is not predicted to vary substantially with CPP, indicating that this signal does not represent an important factor in eliciting the pressure autoregulation response simulated here. This prediction is expected because heart rate does not substantially change with CPP in these experiments. Thus we conclude that sympathetic outflow to vascular smooth muscle also does not substantially change with CPP in these experiments. However, the baseline of the autonomic signal increases from the subepicardium to the subendocardium, indicating that the subendocardium may be more sensitive to changes in sympathetic tone that the other layers.

In all layers the metabolic signal (panel E of Figure 5, 6, and 7) is predicted to make a far greater contribution to the vasoregulation in the hemodilution+dobutamine case compared to the other experimental conditions. In all layers the metabolic signal tends to increase with CPP, and tends to be slightly greater under hemodilution conditions compared to control. The magnitude of the increase with CPP is greatest for the hemodilution+dobutamine case because the oxygen demand is greatest in this condition.

### Model Validation: Simulating Exercise

The ability of the top performing metabolic signal model, *F⸱M*, is validated by evaluating its ability to simulate coronary flow regulation in conscious resting versus exercise conditions. Figure 8 shows input data on aortic and left-ventricular pressures obtained under resting (panel A) and moderate treadmill exercise (2-4 mph, 0% grade, panel B). These pressure data are used to drive the model to match the measured LAD flows illustrated in Figure 9.

**Figure 8.**
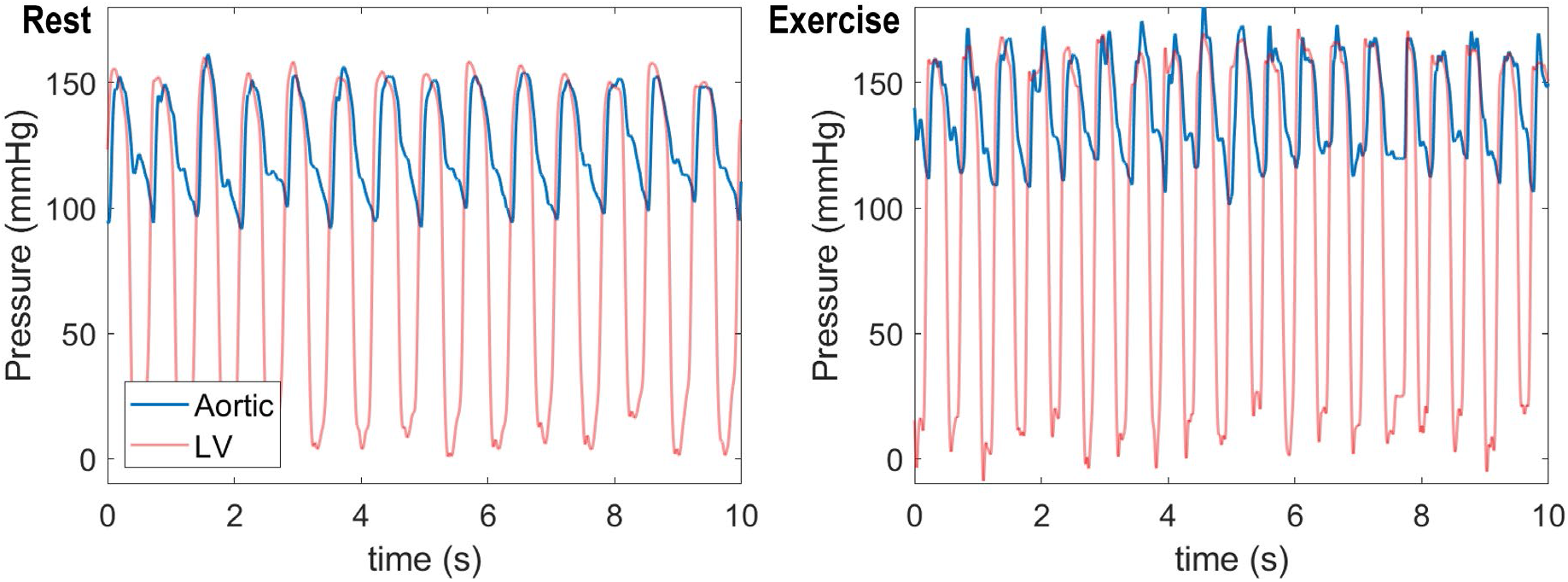
Measured aortic and left-ventricular pressure time courses for a conscious pig in rest (left panel) and exercise (right panel).

**Figure 9.**
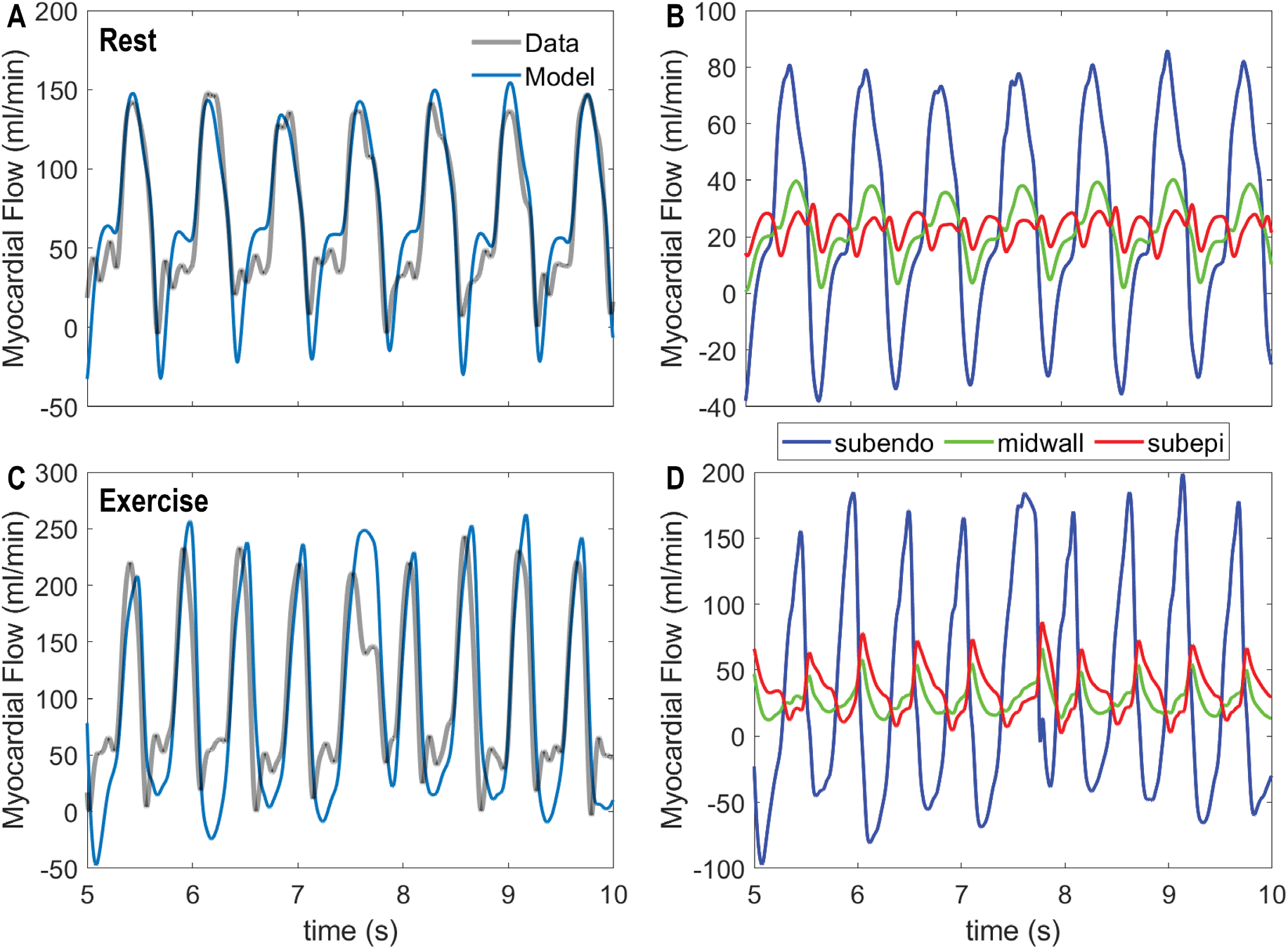
Coronary flow dynamics in rest versus exercise. A. Model-predicted total LAD flow under resting control conditions is compared to measured flow. B. Predicted contributions of subendocardial, midwall, and subepicardial flow to total LAD flow under resting conditions are shown. C. Model-predicted total LAD flow under exercise conditions is compared to measured flow. D. Predicted contributions of subendocardial, midwall, and subepicardial flow to total LAD flow under exercise conditions are shown. In both rest and exercise conditions the model simulations reflect the integrated model using the top-ranked using the ‘*F·M’*, flow times MVO_2_, metabolic signal.

Under resting conditions, the animal’s heart rate was observed to be 84 beats min^−1^, with systolic and diastolic aortic pressures approximately 149 and 102 mmHg, with developed pressure in the ventricle matching peak systolic aortic pressure. During exercise heart rate increases to 114 beats min^−1^ and aortic pressures to 162 and 118 mmHg. We assume a resting average myocardial oxygen consumption rate of 60 μL O_2_/min/g based on Kiel et al. (8). Based on the observed aortic pressures and heart rates, we estimate that left-ventricular mechanical power output increases by 32% in exercise compared to control [24]. And based on the expected linear scaling of power output and oxygen demand, we approximate the exercise oxygen consumption rate to be 79.2 μL O_2_/min/g in exercise.

Given the measured driving pressures and the estimated myocardial oxygen consumption rates, the integrated model can be used to simulate myocardial perfusion in resting and exercise conditions. However, parameters estimated based on the zero-flow pressure experiments (Tables 1 and 2) show a degree of variability and uncertainty. Moreover, they are estimated for different individual animals. Thus, to match the conscious data of Figures 8 and 9, a subset of adjustable parameters was identified, as described in the methods. Adjustable parameters for the conscious experiments are listed in Table 4.

**Table 4.** List of adjustable parameters and estimated values for in vivo simulations

| Parameter | Layer | CoV | Sensitivity | Estimated Value |
| --- | --- | --- | --- | --- |
| $\phi_m$ | Midwall | 0.3084 | 1.38 | 133.577 |
| $C_{myo}$ | Midwall | 0.3075 | 0.13 | 13.923 |
| $C_0$ | Midwall | 0.3405 | 2.07 | -3.296 |
| $\phi_p$ | Subepicardial | 7.6885 | 0.29 | 20.736 |
| $\phi_m$ | Subepicardial | 0.3633 | 2.53 | 75.439 |
| $C_m$ | Subepicardial | 0.5246 | 1.32 | 107.652 |
| $C_{myo}$ | Subepicardial | 0.3010 | 0.28 | 27.483 |
| $C_0$ | Subepicardial | 0.5264 | 0.46 | -9.735 |

Simulated total LAD flow, obtained using the identified F.M model, is compared to measured data for baseline rest (panel A) and exercise (panel C) conditions in Figure 9. The model effectively captures both the pulsatile nature of the flow and the dynamics of flow recruitment in exercise. Panels B and C of Figure 9 show the predicted flow to each of the three myocardial layers, revealing that that the subendocardial flow is positive only during diastole, and becomes negative during systole. In contrast, the subepicardial flow shows positive peaks during both systole and diastole. During exercise, when the driving aortic systolic pressure increases, the systolic peak becomes more important in the subepicardium.

Model predictions of average flow and of the subendocardial-to-subepicardial perfusion ratio are compared to experimental data on Figure 10. In fitting the flow data, the model correctly predicts the transmural variations in perfusion in rest versus exercise. For comparison predictions associated with the best fit of the *M*_*LSv*_ (ranked 5 in Table 3) model are shown in panels A and B. This model is able to match the measured flow, but in doing so predicts subendocardial-to-subepicardial perfusion ratios that are much lower than physiological. These results represent a validation of our identified model and of our model selection procedure. Moreover, the modeling framework provides predictions of transmural constriction/dilation of vessels in terms of their overall vascular tone (e.g., activation *A*), and normalized diameter (Figure 10 C & D). Subendocardial layer tone is predicted to decrease to almost full vasodilation in exercise while the subepicardial layer tone increases. The overall increase in tone in the subepicardial layer is overcome by the pressure-induced dilation (panel D).

**Figure 10.**
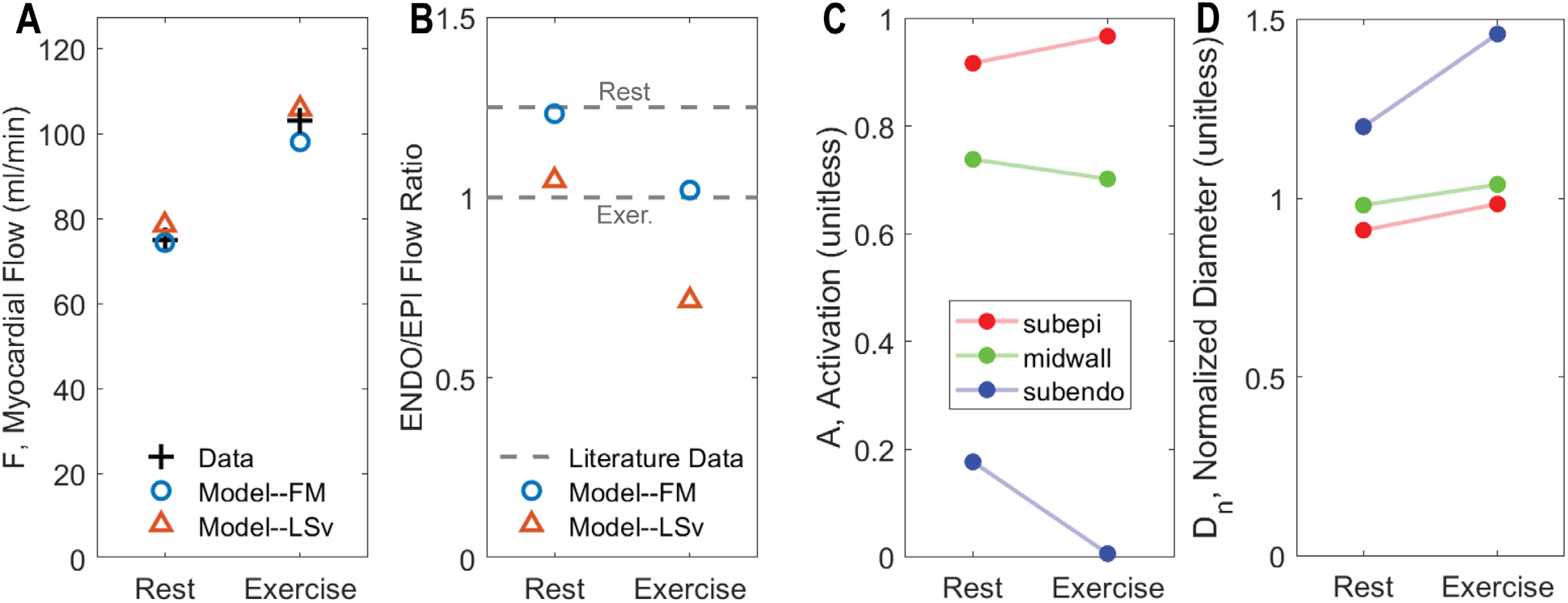
Regulation of coronary flow in exercise. A. Comparison of measured to model-predicted total LAD flow in resting versus exercise condition. B. Comparison of measured to model-predicted subendocardial-to-subepicardial (ENDO/EPI) flow in resting versus exercise condition. In panels A and B fits from competing metabolic model are illustrated, showing that the lower ranking model fails to capture the ENDO/EPI flow ratio. C. Predicted total vascular tone activation in each layer of the myocardium for resting and exercise conditions. D. Predicted representative resistance vessel diameter in each layer of the myocardium for resting and exercise conditions.

Figure 11 illustrates how each regulatory signal contributes to the overall predicted response to exercise. The figures plots the exercise-induced change in each stimuli 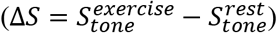 in each layer of the myocardium for the F.M model. The model-predicted vasoconstriction in the subepicardium in exercise is shown to be due to a myogenic response to elevated perfusion pressure. The autonomic vasodilation signal (*S*_*HR*_) is predicted to preferentially influences the subendocardium, with a negligible on the subepicardium.

**Figure 11.**
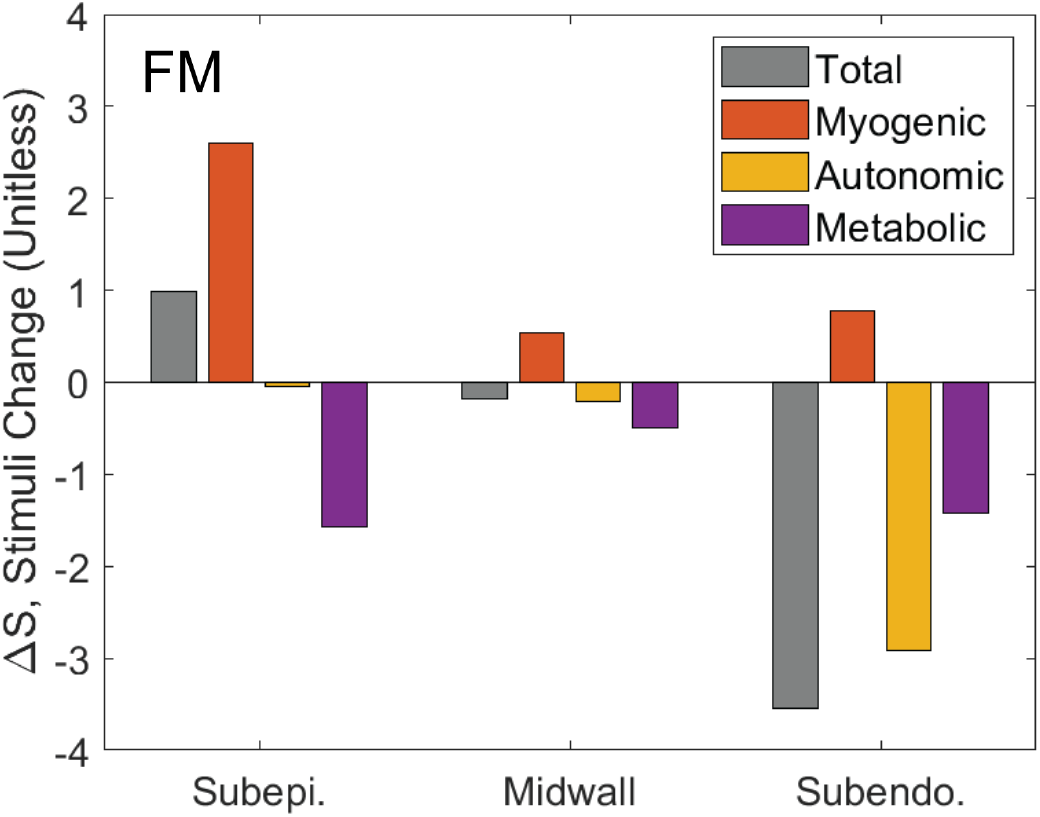
The change in total, myogenic, metabolic, and autonomic stimuli in response to exercise. The gray bars show the net total change in stimulus in each layer exercising compared to rest conditions. The individual contributors to the stimuli signal—myogenic, autonomic, and metabolic—sum to the total in each layer. The total stimulus in the subepicardium is higher in exercise than in rest, resulting in an increased activation. The primary contributor to the increased activation in the subepicardium in exercise is the myogenic stimulus. In the subepicardium exercise causes an overall reduction in vascular tone, caused by a combination of metabolic and autonomic stimuli.

## Discussion

The physiological balance between coronary blood and myocardial oxygen consumption (MVO_2_) is preserved over a variety of physiological and pathophysiologic condition, including in the response to exercise where increases in oxygen demand are met by corresponding increases in oxygen delivery.

### Modeling framework and the overall approach

This study integrates a lumped approach to simulating ventricular-vascular interaction and myocardial perfusion adapted from Spaan et al. [9], and Mynard et al. [5, 10], a vasoregulation model adapted from Carlson et al. [6], and a panel of competing mechanisms representing metabolic control of myocardial blood flow. The multi-scale modeling framework is illustrated in Figure 2. The model components are identified based on analysis of data from zero-flow pressure experiments, as illustrated in Figure 1. Based on their relative ability to match the experimental data, ranked based on goodness of fit and degrees of freedom in the models, the maximally likely metabolic mechanism is identified, and further validated based on simulations of conscious response to exercise (Figures 8, 9, and 10). One of the major novelties of our study is the application of established tools for model discrimination in the context of multi-scale systems with high degrees of uncertainly and structural and parametric freedom.

### Identification of hypothetical metabolic signal

Data from three conditions and four experimental animals were analyzed with model formulations that integrated seven competing hypotheses for the nature of the metabolic feedback signal. The competing hypotheses are summarized in Table 3, along with a summary ranking of most-to-least likely mechanism. These results illustrate a combined structural and parametric model identification, with several competing models ruled out as not viable. In the top-ranked model, the metabolic signal is represented as proportional to metabolic rate and flow. Our preliminary interpretation of this result is that it represents an ephemeral metabolic signal for which production is proportional to MVO_2_ and delivery proportional to flow. Potential candidates for such a signal include short-lived reactive species such as H_2_O_2_ [7].

Predicted resistance vessel diameters are plotted as functions of transmural pressure at different CPP values for the midwall layer in one experimental animal in Figure 4. Predicted diameters are plotted as functions of CPP for all animals under all conditions in Supplementary Figures S14-S25. Simulations consistently show a greater resistance vessel diameter for the hemodilution compared to control case. These differences are small but important because they are consistently required to match the data on flow and pressure. In fact, the reason why the ATP-mediated metabolic model (Metabolic Signal #1, Table 3) cannot effectively match data from the zero-flow pressure experiment is that it cannot capture this phenomenon. In other words, even with arbitrarily adjusting parameters, the output of the model capturing the ATP hypothesis cannot be made to be consistent the observed pressure response to upstream occlusion. Therefore, by combining the lumped microcirculation model, a simple vessel mechanics model, and our ATP-mediated metabolic control model to analyze these data, our analysis argues against the hypothesis that ATP derived from red blood cells represents the primary metabolic signal for coronary blood flow regulation. Rigorously, discarded models are judged as non-viable only in the form that they are implemented for this analysis. Nevertheless, this finding highlights the sort of novel insight that comes from analyzing time-dependent data using even a relatively crude multi-scale and spatially distributed modeling framework.

Thus although we do not interpret the top-ranked model as “proven” or “correct”, the disproof of the alternatives usefully narrows the scope of possibilities. Furthermore, even if the basic formulation of the F·M model correctly represents the basic action of metabolic mechanism, this analysis does not identify the molecular mechanism underlying the putative mechanism. Rather, we interpret the ephemeral metabolic signal F·M model as a working hypothesis to test and refine with further experiments and models that represent the metabolic and signaling mechanisms with more mechanistic resolution. The validity of this working hypothesis is supported by the ability of the associated model to simulate the response in myocardial perfusion to exercise.

### Matching demand in exercise

The ability of the ephemeral signal hypothesis, when integrated in this modeling framework, to effectively simulate the dynamics of myocardial perfusion in the transition from resting state to exercise is illustrated in Figures 8, 9, and 10. Because lower-ranked models (e.g., Metabolic Signal #5) predict a greater than physiological drop in subendocardial perfusion with exercise, these results represent a validation of the maximal-likelihood evaluation process: the top-ranked model captures behavior not used for identification, while lower ranked models fail.

Simulation of the exercise response also yields novel hypotheses regarding the regional response to increasing exercise demand. For example, simulations predict that subepicardial vascular tone is higher during exercise than at rest, due to a myogenic response to elevated perfusion pressure. Another prediction associated with the exercise simulations is that sympathetic-mediated vasodilation preferentially influences the subendocardium. The physiological role of sympathetic activation in the control of myocardial perfusion is much more complex than the simplified way it is treated in the current model, which does not take into account a potentially heterogeneous distribution of α-mediated constriction effectors [2, 25]. Moreover, measurement of β_2_-adrenergic receptor densities show transmurally uniform expression in the human myocardium [26, 27]. Furthermore, subepicardial resistance arterioles from pig have been observed to show a greater vasodilation response to selective β_2_ stimulation and have greater β_2_ receptor density than subendocardial resistance arterioles [28]. At the same time, α-mediated vasoconstriction has been shown to preferentially distribute flow to the subendocardium, particularly during exercise and in the presence of a proximal stenosis [29]. Our model lumps α- and β-mediated effects on resistance vessels into a single signal, *S*_*HR*_. Thus the model prediction that the gain associated with this signal is greater in the subendo- compared to subepicardium is consistent with the interpretation that the overall response to sympathetic stimulation is relatively more vasodilatory in the subendocardium compared to the subepicardium. Moreover, this overall transmural gradient in the response of vascular tone in exercise is undoubtedly influenced by regional heterogeneities in responses of vessels of different sizes and transmural depth to metabolic, myogenic, and autonomic signals. A deeper investigation of how these factors work together in vivo may be possible by combining models with more anatomically resolved detail [30] with the regulatory mechanisms analyzed here.

### Limitations of the study

The current model uses a simplified lumped-parameter of myocardial circulation downstream of the LAD, capturing regional differences in perfusion only in terms of a relatively coarse representation of transmural heterogeneity in the LV free wall. Previous applications of this approach have coupled the myocardial perfusion model to a 1D hemodynamics models of epicardial vessels to simulate regional perfusion, for example in the left-ventricular free wall versus the septum [5, 10]. Moreover, the target transmural flow distribution (ENDO/EPI ratio) in the current study is estimated literature reports, which consistently report ENDO/EPI at only one experimental point (control, CPP=100 mmHg). More data on variation of transmural perfusion with CPP would be valuable in constraining and testing model behavior. More broadly, a rigorous uncertainty quantification will be useful in analyzing measurement errors propagated to the final model predictions. Finally, the current model assumes the intramyocardial pressure is a linear function of left ventricular pressure in all cases. This assumption has to be re-examined especially for dobutamine experiments which involve elevations in the myocardial contractility which directly influences the intramyocardial pressure via changes in the myocyte shortening induced pressure [31].

## Model Code Availability

Computer codes, implemented in the MATLAB computing environment, are available for download from the repository at https://github.com/beards-lab/Multiscale-Coronary-Flow-Control.

## References

1. Dole, W.P. and D.W. Nuno, Myocardial oxygen tension determines the degree and pressure range of coronary autoregulation. Circ Res, 1986. 59(2): 202–15.

2. Duncker, D.J. and R.J. Bache, Regulation of coronary blood flow during exercise. Physiol Rev, 2008. 88(3): 1009–86.

3. Tune, J.D., et al., Disentangling the Gordian knot of local metabolic control of coronary blood flow. Am J Physiol Heart Circ Physiol, 2020. 318(1): H11–H24.

4. Kiel, A.M., et al., Local metabolic hypothesis is not sufficient to explain coronary autoregulatory behavior. Basic Res Cardiol, 2018. 113(5): 33.

5. Mynard, J.P., D.J. Penny, and J.J. Smolich, Scalability and in vivo validation of a multiscale numerical model of the left coronary circulation. Am J Physiol Heart Circ Physiol, 2014. 306(4): H517–28.

6. Carlson, B.E. and T.W. Secomb, A theoretical model for the myogenic response based on the length-tension characteristics of vascular smooth muscle. Microcirculation, 2005. 12(4): 327–38.

7. Saitoh, S., et al., Hydrogen peroxide: a feed-forward dilator that couples myocardial metabolism to coronary blood flow. Arterioscler Thromb Vasc Biol, 2006. 26(12): 2614–21.

8. Berwick, Z.C., et al., Contribution of voltage-dependent K(+) channels to metabolic control of coronary blood flow. J Mol Cell Cardiol, 2012. 52(4): 912–9.

9. Spaan, J.A., et al., Dynamics of flow, resistance, and intramural vascular volume in canine coronary circulation. Am J Physiol Heart Circ Physiol, 2000. 278(2): H383–403.

10. Mynard, J.P. and J.J. Smolich, Influence of anatomical dominance and hypertension on coronary conduit arterial and microcirculatory flow patterns: a multiscale modeling study. Am J Physiol Heart Circ Physiol, 2016. 311(1): H11–23.

11. Ho, S.Y., D. Sanchez-Quintana, and A.E. Becker, A review of the coronary venous system: a road less travelled. Heart Rhythm, 2004. 1(1): 107–12.

12. Hoffman, J.I. and J.A. Spaan, Pressure-flow relations in coronary circulation. Physiol Rev, 1990. 70(2): 331–90.

13. Breisch, E.A., et al., Cardiac vasculature and flow during pressure-overload hypertrophy. Am J Physiol, 1986. 251(5 Pt 2): H1031–7.

14. Breisch, E.A., et al., Exercise-induced cardiac hypertrophy: a correlation of blood flow and microvasculature. J Appl Physiol (1985), 1986. 60(4): 1259–67.

15. Krombach, R.S., et al., Amlodipine therapy in congestive heart failure: hemodynamic and neurohormonal effects at rest and after treadmill exercise. Am J Cardiol, 1999. 84(4A): 3L–15L.

16. Carlson, B.E., J.C. Arciero, and T.W. Secomb, Theoretical model of blood flow autoregulation: roles of myogenic, shear-dependent, and metabolic responses. Am J Physiol Heart Circ Physiol, 2008. 295(4): H1572–9.

17. Young, J.M., et al., Slackness between vessel and myocardium is necessary for coronary flow reserve. Am J Physiol Heart Circ Physiol, 2012. 302(11): H2230–42.

18. Pradhan, R.K., et al., Open-loop (feed-forward) and feedback control of coronary blood flow during exercise, cardiac pacing, and pressure changes. Am J Physiol Heart Circ Physiol, 2016. 310(11): H1683–94.

19. Holtz, J., et al., Intracapillary hemoglobin oxygen saturation and oxygen consumption in different layers of the left ventricular myocardium. Pflugers Arch, 1977. 370(3): 253–8.

20. Vinten-Johansen, J. and H.R. Weiss, Oxygen consumption in subepicardial and subendocardial regions of the canine left ventricle. The effect of experimental acute valvular aortic stenosis. Circ Res, 1980. 46(1): 139–45.

21. Snyder, G.K., Influence of temperature and hematocrit on blood viscosity. Am J Physiol, 1971. 220(6): 1667–72.

22. Navakatikyan, M.A., A model for residence time in concurrent variable interval performance. J Exp Anal Behav, 2007. 87(1): 121–41.

23. Vinnakota, K.C., et al., Analysis of the diffusion of Ras2 in Saccharomyces cerevisiae using fluorescence recovery after photobleaching. Phys Biol, 2010. 7(2): 026011.

24. Holmberg, S., W. Serzysko, and E. Varnauskas, Coronary circulation during heavy exercise in control subjects and patients with coronary heart disease. Acta Med Scand, 1971. 190(6): 465–80.

25. Goodwill, A.G., et al., Regulation of Coronary Blood Flow. Compr Physiol, 2017. 7(2): 321–382.

26. Murphree, S.S. and J.E. Saffitz, Distribution of beta-adrenergic receptors in failing human myocardium. Implications for mechanisms of down-regulation. Circulation, 1989. 79(6): 1214–25.

27. Beau, S.L., T.K. Tolley, and J.E. Saffitz, Heterogeneous transmural distribution of beta-adrenergic receptor subtypes in failing human hearts. Circulation, 1993. 88(6): 2501–9.

28. Hein, T.W., et al., Heterogeneous beta2-adrenoceptor expression and dilation in coronary arterioles across the left ventricular wall. Circulation, 2004. 110(17): 2708–12.

29. Chilian, W.M. and P.H. Ackell, Transmural differences in sympathetic coronary constriction during exercise in the presence of coronary stenosis. Circ Res, 1988. 62(2): 216–25.

30. Namani, R., et al., Effects of myocardial function and systemic circulation on regional coronary perfusion. J Appl Physiol (1985), 2020. 128(5): 1106–1122.

31. Algranati, D., G.S. Kassab, and Y. Lanir, Mechanisms of myocardium-coronary vessel interaction. Am J Physiol Heart Circ Physiol, 2010. 298(3): H861–73.

