## Supplementary Figures for "Multiscale model of the physiological control of myocardial perfusion to delineate putative metabolic feedback mechanisms"

#### S1. Aortic and left ventricular pressure

The aortic pressure ($P_{ao}$) is continuously measured during the zero-flow pressure measurements (before and after occlusion). To estimate left ventricular pressure $P_{lv}$, half-sine functions were used to match AoP in the systolic phase. Figure S1 shows $P_{ao}$ measurements and estimated $P_{lv}$ for Pig D (control) experiment with 100 mmHg CPP. Analogous data exist for all experiments.

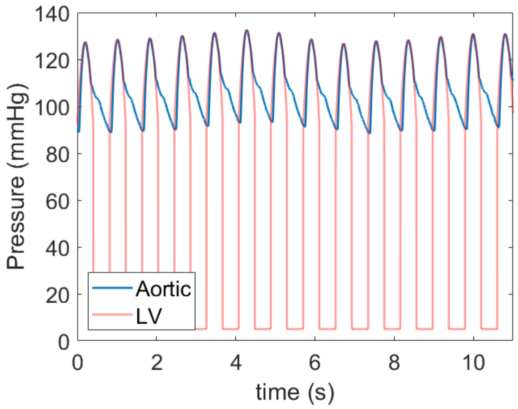


Figure S1. Measured aortic and estimated left ventricular pressure for Pig D, CPP = 100 mmHg.

#### S2. Myocardial circulation model fits

Model fits to zero-flow pressure time course data for all animals/experimental conditions are illustrated below.

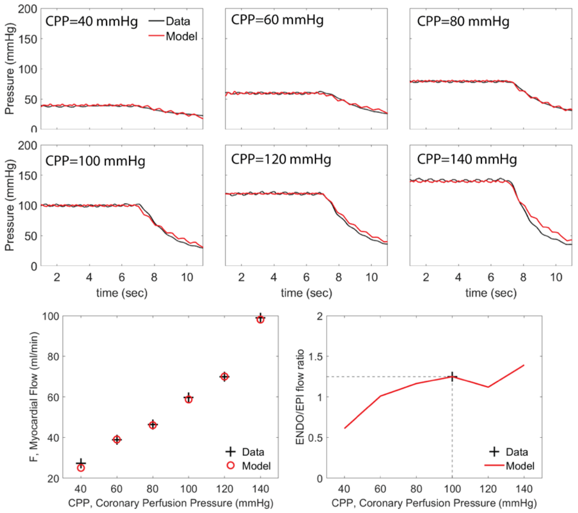


Figure S2. Data and model fits for Pig A under control conditions.

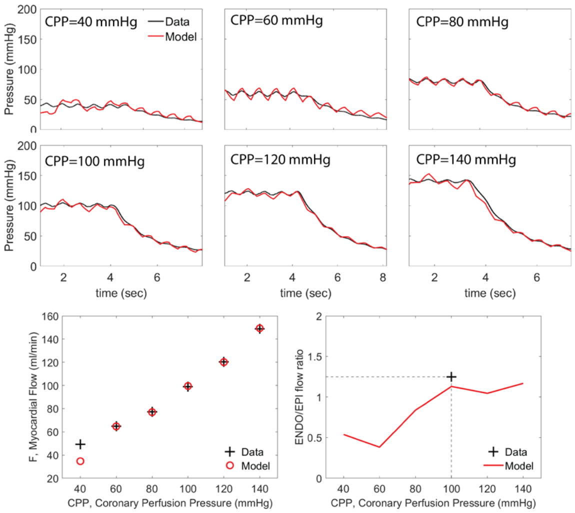


Figure S3. Data and model fits for Pig A under hemodilution conditions.

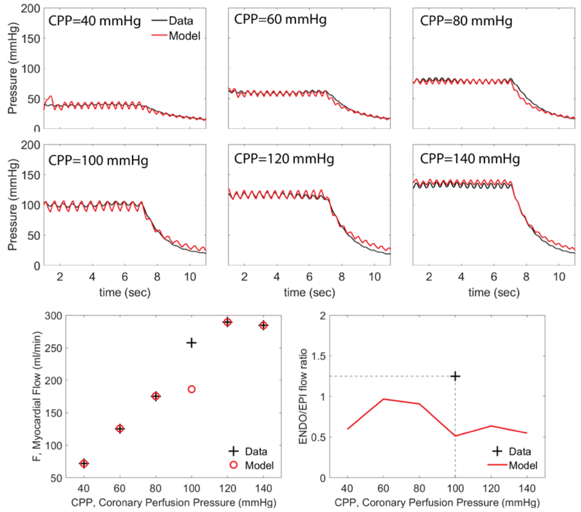


Figure S4. Data and model fits for Pig A under hemodilution+dobutamine conditions.

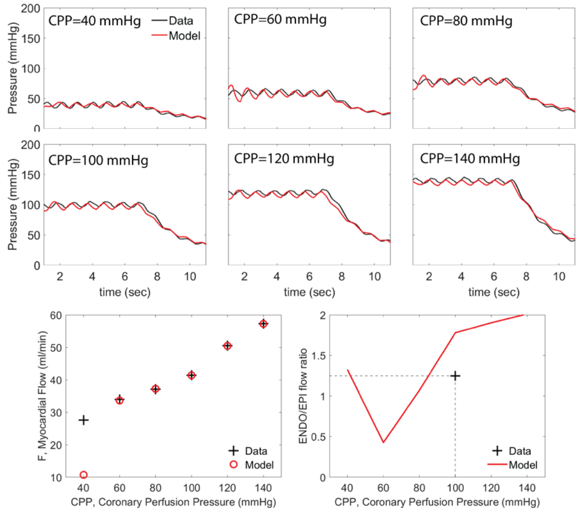


Figure S5. Data and model fits for Pig B under control conditions.

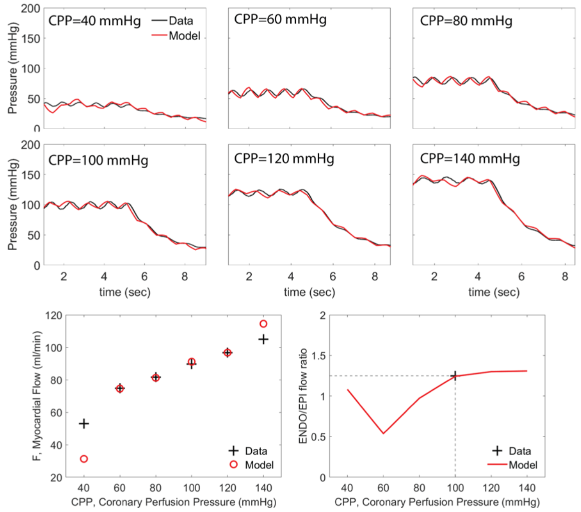


Figure S6. Data and model fits for Pig B under hemodilution conditions.

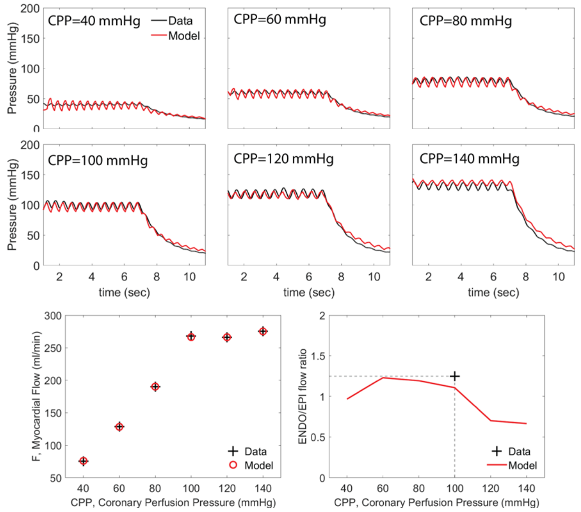


Figure S7. Data and model fits for Pig B under hemodilution+dobutamine conditions.

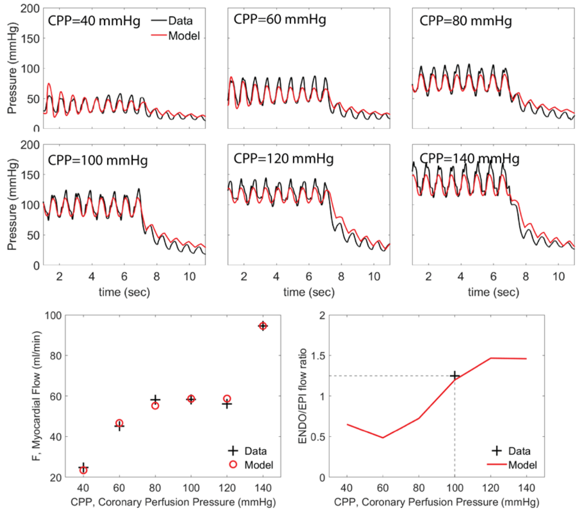


Figure S8. Data and model fits for Pig C under control conditions.

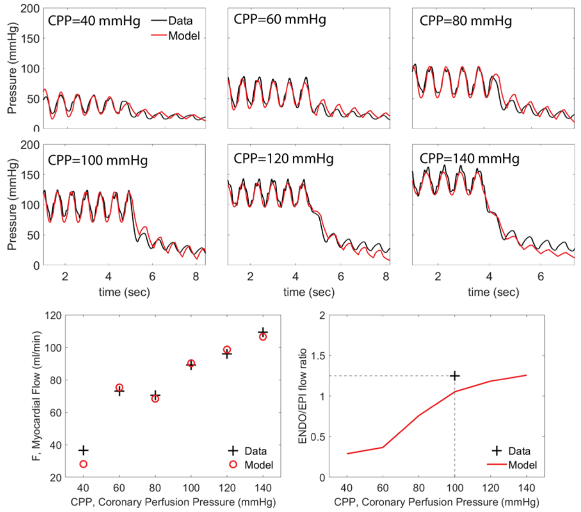


Figure S9. Data and model fits for Pig C under hemodilution conditions.

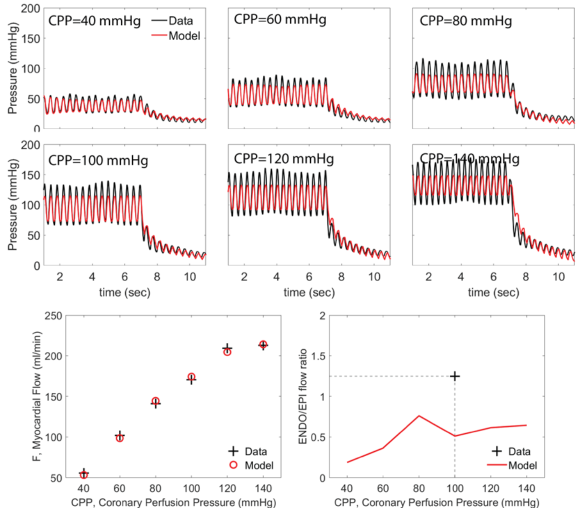


Figure S10. Data and model fits for Pig C under hemodilution+dobutamine conditions.

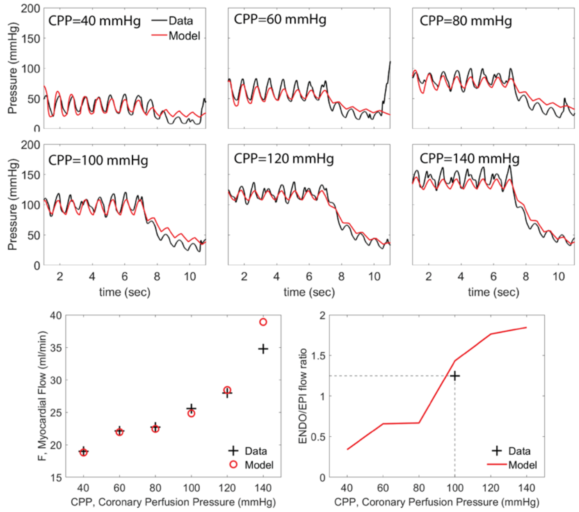


Figure S11. Data and model fits for Pig D under control conditions.

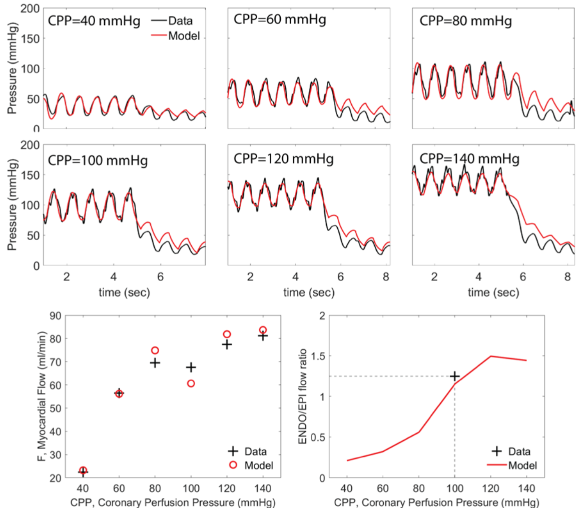


Figure S12. Data and model fits for Pig D under hemodilution conditions.

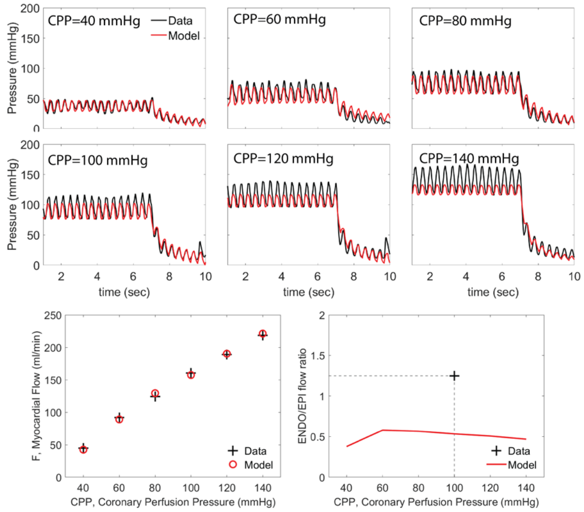


Figure S13. Data and model fits for Pig D under hemodilution+dobutamine conditions.

### S3. Representative vessel model fits

The representative vessel model fits with the top-ranking metabolic signal $MS_{FM}$ are illustrated below. Each figure plots: Panel A: equivalent diameters as functions of transmural pressure in the representative vessel, with markers “+” and “o” showing Model 1 and Model 2 results, respectively; Panel B: predicted vessel activation vs. transmural pressure; Panels C-E: predicted regulatory signals strengths as functions transmural pressure.

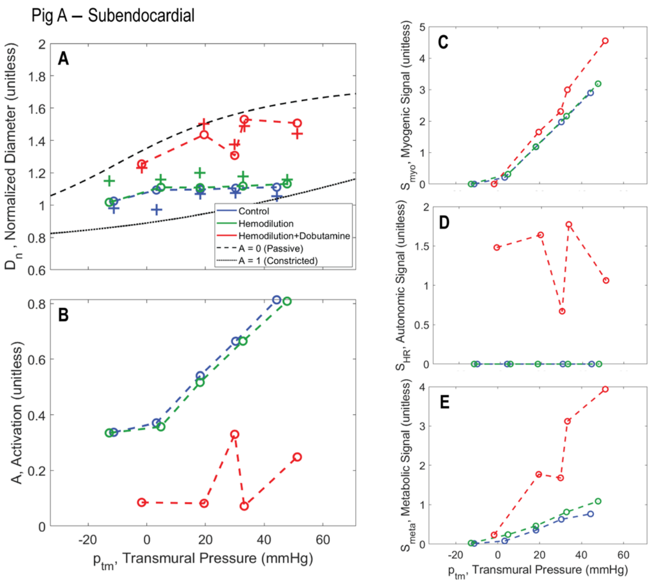


Figure S14.

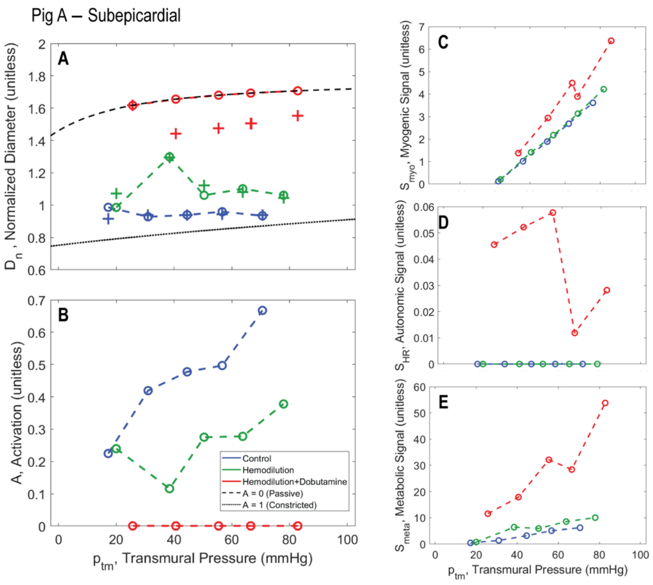


Figure S16.

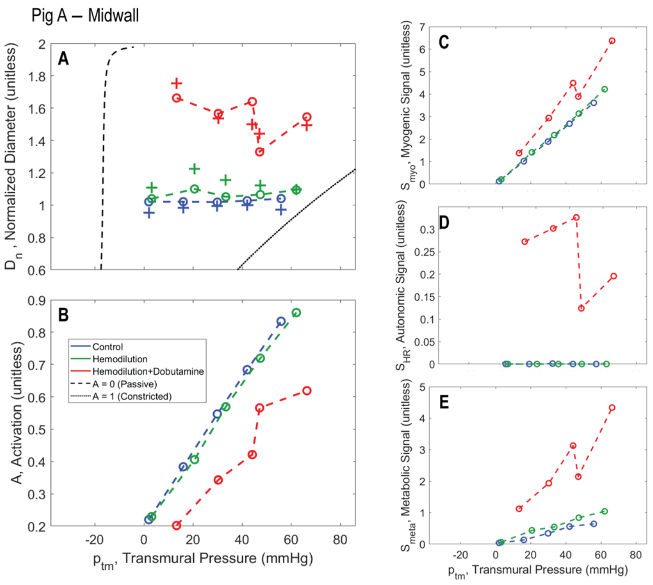


Figure S15.

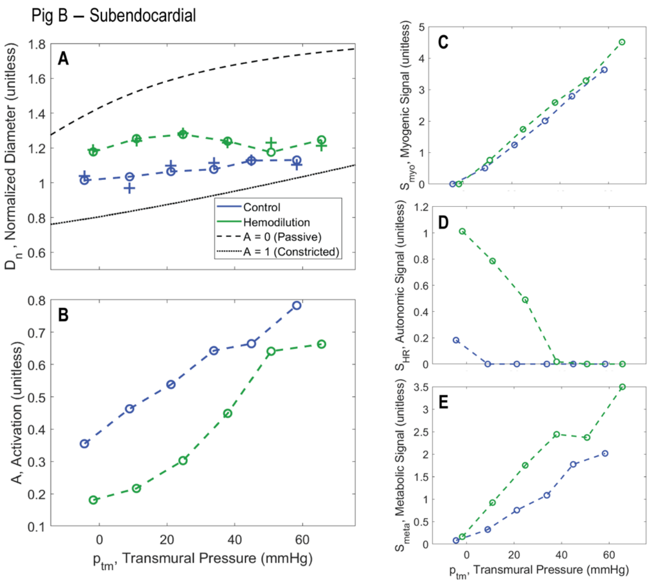


Figure S17.

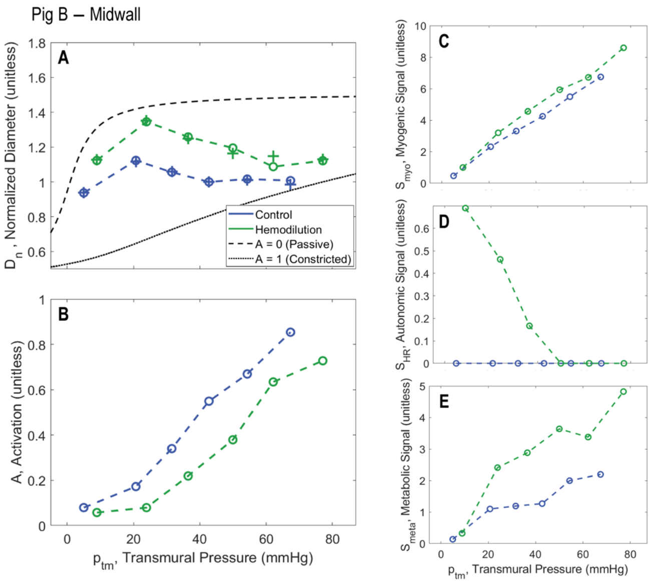


Figure S18.

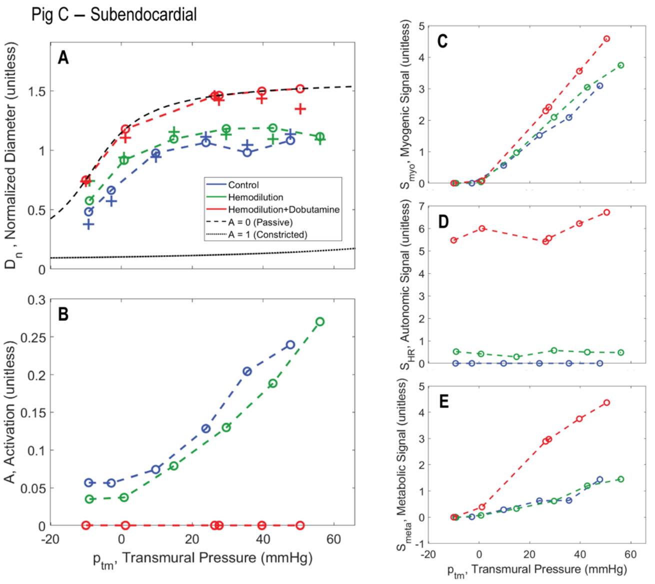


Figure S20.

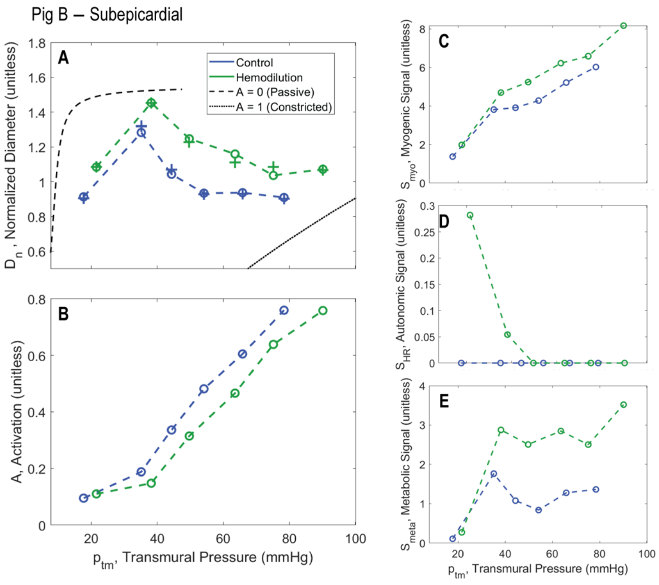


Figure S19.

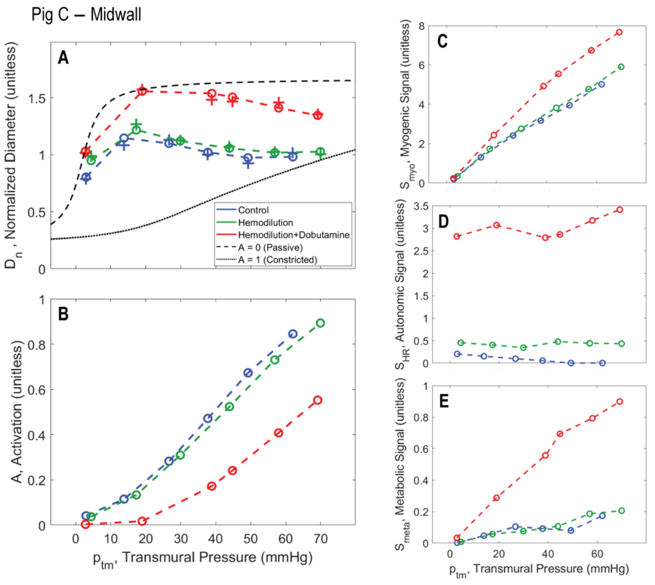


Figure S21.

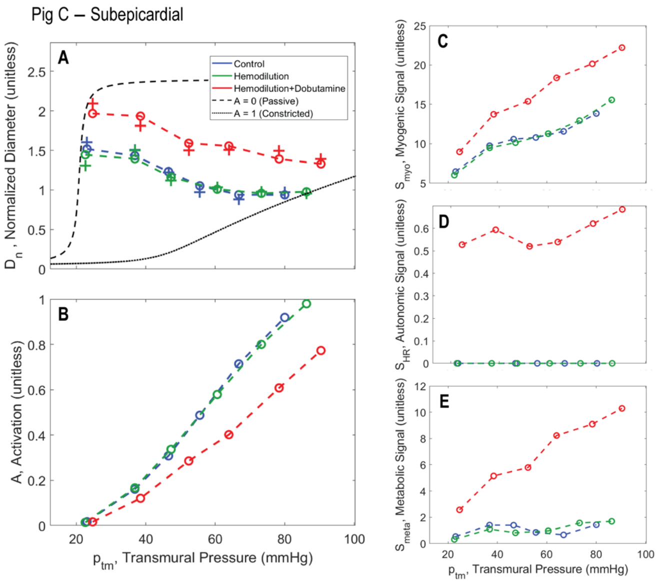


Figure S22.

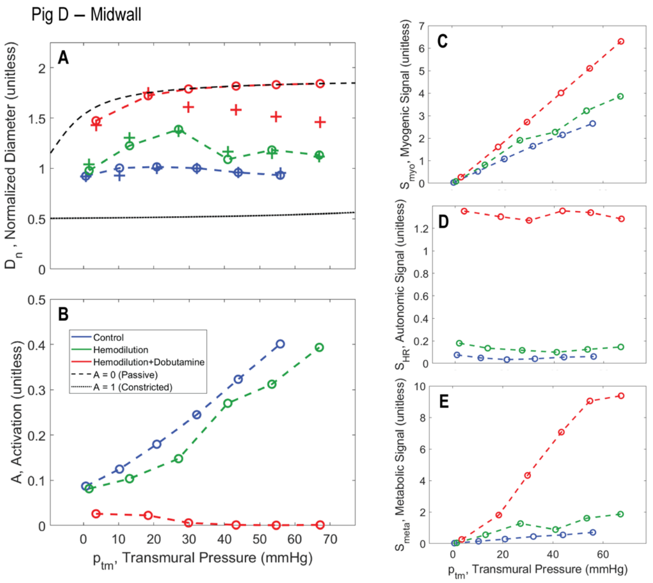


Figure S24.

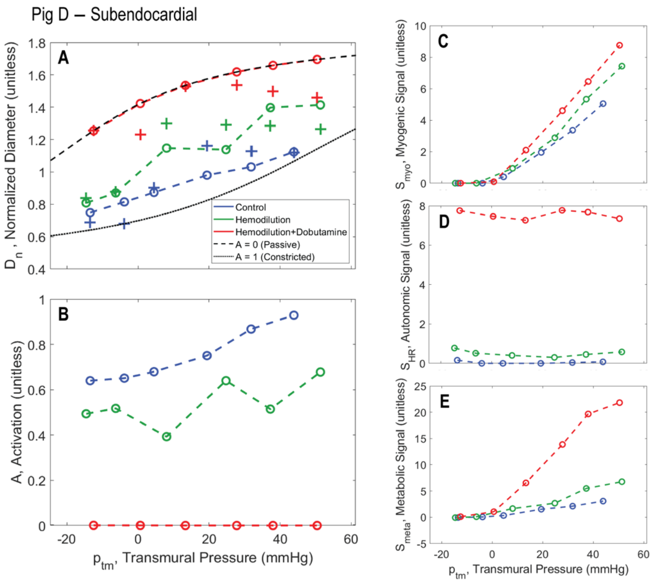


Figure S23.

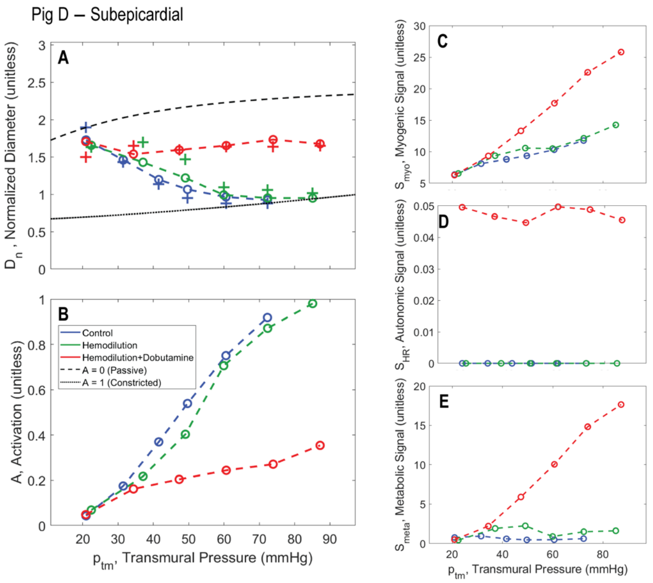


Figure S25.
